# After-hyperpolarization promotes the firing of mitral cells through a voltage dependent modification of action potential threshold

**DOI:** 10.1101/2021.04.23.441072

**Authors:** Fourcaud-Trocmé Nicolas, Zbili Mickaël, Duchamp-Viret Patricia, Kuczewski Nicola

## Abstract

In the olfactory bulb (OB), mitral cells (MCs) display a spontaneous firing that is characterized by bursts of action potentials intermixed with silent periods. Burst firing frequency and duration are heterogeneous among MCs and increase with membrane depolarization. By using patch clamp recording on rat slices, we dissected out the intrinsic properties responsible of this activity. We showed that the threshold of action potential (AP) generation dynamically changes as a function of the trajectory of the membrane potential; becoming more negative when the membrane was hyperpolarized and having a recovering rate, inversely proportional to the membrane repolarization rate. Such variations appeared to be produced by changes in the inactivation state of voltage dependent Na^+^ channels. Thus, the modification AP threshold favours the initiation of the burst following hyperpolarizing event such as negative membrane oscillations or inhibitory transmission. After the first AP, the following afterhyperpolarization (AHP) brought the threshold just below the membrane resting potential or within membrane oscillations and, as a consequence, the threshold was exceeded during the fast repolarization component of the AHP. In this way the fast AHP acts as a regenerative mechanism that sustains the firing. Bursts were stopped by the development of a slow repolarization component of the AHP. The AHP characteristics appeared as determining the bursting properties; AHP with larger amplitudes and faster repolarizations being associated with longer and higher frequency bursts. Thus, the increase of bursts length and frequency upon membrane depolarization would be attributable to the modifications of the AHP and of Na^+^ channels inactivation.

## Introduction

The after-hyperpolarization (AHP) that follows the action potential (AP) is generally seen as an inhibitory mechanism that limits the neuronal activity, by promoting firing frequency adaptation and termination of AP burst (Schwindt *et al.,* 1988; Faber & Sah, 2007; Adelman *et al.,* 2012; Reuveni & Barkai, 2018). The main mechanism underlying AHP is the activation of voltage- and calcium-dependent potassium currents. However, in some neuronal types, such as MCs, main output neurons of the OB, the synaptic transmission also contributes to AHP shape (Duménieu *et al.,* 2015). Differences in activation-inactivation kinetics and calcium sensitivity of the different subtypes of potassium channels, underlying AHP, are responsible for a splitting of its course into three successive components, a fast (fAHP), a medium (mAHP) and a slow (sAHP), that differ in onset times, rise and decay kinetics (Schwindt *et al.,* 1988; Sah & Faber, 2002; Andrade *et al.,* 2012). The relative contribution of each components evolves during the neuronal discharge making the AHP shape dependent on preceding neuronal activity (Duménieu *et al.*, 2015). The inhibitory action of the AHP is generally attributed to the potassium channels which prevent excitatory currents to bring the membrane potential (MP) to the AP threshold (Rubin & Cleland, 2006). This vision may however lead to neglect the possibility that AHP could, in addition, result in the deinactivation of some voltage dependent channels, such as calcium T-type and sodium ones (Deister *et al.,* 2009; Cain & Snutch, 2010; Platkiewicz & Brette, 2011; Iyer *et al.,* 2017). In this way, such a deinactivation would promote neuronal firing when the MP moves back to the resting state value. In OB, MCs present a spontaneous firing activity that is characterized by AP clusters interspaced by silent periods (Desmaisons *et al.,* 1999). This activity is mainly due to intrinsic membrane properties, since it was observed in pharmacologically isolated MCs, in OB slices (Balu *et al.,* 2004). The cellular mechanisms behind bursting activity of MCs remain to be elucidated. Balu and Strowbridge (2004) proposed that burst of APs ends up thanks to the build-up of the slow AHP while a computational model suggested (Rubin & Cleland, 2006) that burst termination would rather rely on the accumulation of potassium I_A_ current during the firing. However, the mechanisms that trigger the burst, maintain the sustained firing, determine the firing properties and the number of APs or their frequency, remain largely unknown. Here we provide evidences that, through a dynamic change of AP threshold, the AHP-induced modification of the MP would act both as a burst regenerative process as well as a mechanism of burst termination. In this way the AHP features would play a pivotal role in determining the firing properties of MCs.

## Methods

### Animals

Animal handling was conducted in accordance with the European Community Council Directive 86/609/EEC. Experiments were performed in P30-P42 male Long Evans rats (Janvier, Le Genest-Saint-Isle, France). The animals were maintained on a normal light cycle and ad libitum accessed to water and food.

### Slice preparation

Animals were anaesthetized with an intra-peritoneal injection of ketamine (50 mg/ml) and then, decapitated. The head was quickly immersed in ice-cold (2-4°C) carbogenized artificial cerebrospinal fluid (ACSF; composition: 125 mM NaCl, 4 mM KCl, 25 mM NaHCO3, 0.5 mM CaCl2, 1.25 mM NaH2PO4, 7 mM MgCl2 and 5.5 mM glucose; pH = 7.4) oxygenated with 95% O2, 5% CO2. The osmolarity was adjusted to 320 mOsm with sucrose. The two olfactory bulbs (OBs) were removed from the cranial cavity and cut in horizontal slices (400 μm thick) using a Leica VT1000s vibratome (Leica Biosystems, France). Slices were then incubated in Gibb’s chamber at 30 ±1°C in modified calcium and magnesium ACSF (CaCl2=2 mM and MgCl2 = 1 mM).

### Electrophysiological recordings

Slices were transferred into a recording chamber mounted on an upright microscope (Axioskop FS, Zeiss) and perfused with oxygenated ACSF (4 ml/min) at 30 ±1°C. Neurons were visualized using a 40X objective and a Hamamatsu “Orca Flash 4.0” camera. Measurements were performed with a RK 400 amplifier (BioLogic, France). Data were acquired with a sampling frequency of 25 kHz on a PC-Pentium D computer using a 12-bit A/D-D/A converter (Digidata 1440A, Axon Instruments) and PClamp10 software (Axon Instruments). Patch-clamp recordings were achieved with borosilicate pipettes (o.d.: 1.5 mm; i.d.: 1.17 mm; Clark Electromedical Instruments), filled with the intracellular solution (131 mM K-gluconate, 10 mM HEPES, 1 mM EGTA, 1 mM MgCl2, 2 mM ATP-Na2, 0.3 mM GTP-Na3, and 10 mM phosphocreatine; pH = 7.3, 290 mOsm). In our experimental conditions, the equilibrium potential of chloride ions (ECl) was −110 mV, and that of potassium ions (Ek) was −92 mV. The calculated junction potential of 13 mV was corrected offline.

### Data analysis

#### Evoked activity

Experiments were performed in current clamp. A small steady membrane hyperpolarization was ensured by negative current injection in order to prevent spontaneous firing. For the experiments investigating the relationship between the level of hyperpolarization and the AP threshold, two APs were generated by two 3ms depolarizing current steps applied at 6s interval; the second step being preceded by membrane hyperpolarization varying in amplitude and duration. Some data were excluded from the analysis when the average Vrest, in the 500 ms preceding the depolarizing step, differed by more than 2 mV between the two evoked APs. In experiments investigating the relationship between the speed of repolarization-hyperpolarization and AP threshold, current ramps having variable slopes were applied. The AP threshold was calculated from the first AP generated during the ramp: it was defined as being the first point with a strict positive acceleration (second derivative of its MP) during the AP rising phase, before it reaches its maximum depolarizing rate.

#### Sodium currents

Experiment were performed in voltage clamp in the presence of 0.3 mM cadmium, 4 mM Nickel 4, 10 mM Tetraethylammonium (TEA), 10 mM 4-Aminopyridine (4AP), 10 mM 2,3-Dioxo-6-nitro-1,2,3,4-tetrahydrobenzo[f]quinoxaline-7-sulfonamide disodium salt (NBQX), 5 mM 2-APV, D-APV, D-2-amino-5-phosphonovalerate (D-APV) and 5 Mm Bicuculline.

#### Spontaneous activity

APs were detected as membrane depolarizations crossing −23mV, and the minimum interspike interval was set at 1ms. In one cell, the crossing threshold was set at −43mV because of the low amplitude of first AP in bursts.

Burst were detected based on a maximum threshold for the interspike intervals (tISI) below which, all occurring APs were assigned to the same burst. To achieve this, tISI was first set at 90 ms for all analysed MCs, and then adapted cell by cell in a recursive manner through the following procedure: we computed a new tISI as the median ISI of all detected bursts in a cell, plus 4 times their median absolute deviation (except for one cell: only 0.8 times its median absolute deviation, cell label 8 in the figures). If the new tISI was lower than the previous one, we used it to detect again bursts of this cell and restarted over a new tISI computation and so on, until the new tISI was larger than the last one (final burst ISI thresholds: mean: 47ms, SD: 19ms, range 13-90ms). The reliability of the burst detection method was assessed by visual inspection of traces.

Because some recorded MCs showed slow fluctuations of subthreshold MP, the latter was computed before each burst. If the interval without AP preceding a burst (interburst interval) lasted at least 200ms, we defined the burst resting potential (Vrest) as the median MP during the interburst interval (excluding half of the tISI at the beginning and 5ms at the end). We also defined subthreshold fluctuation amplitude as the maximum MP on the same interval. If the interburst interval lasted less than 200ms, its Vrest and subthreshold fluctuation amplitudes were defined as the same as for the preceding burst. In some figures (see legends), traces are aligned on Vrest which was then arbitrary set at 0mV.

AP threshold was defined as the first point with a strictly positive acceleration (second derivative of MP) during the AP rising phase before MP reaches its maximum positive acceleration rate. In this study, we often used the relative AP threshold defined as the difference between AP threshold and Vrest.

The AP prepotential was the first negative MP peak that preceded the AP threshold.

It was detected by stepping backward in time from AP threshold by 1ms steps and, stopping as soon as a MP rebound of at least 0.4mV (relatively to the lowest MP in the interval from current time step up to AP threshold time) was found. The AP prepotential was defined as the lowest MP between the rebound time and the AP threshold time.

Pre-AP slope was the slope of the MP course, preceding the first AP of the burst. When AP prepotential occurred more than 5ms before the AP threshold time, the pre-AP slope was obtained by a linear regression of the MP course during the 5ms before AP threshold. When AP prepotential occurred less than 5ms before AP threshold, the pre-AP slope was obtained by a linear regression of MP course in a range of 20% to 80% from the AP-prepotential (0%) to the AP threshold (100%).

AHP amplitude was calculated by making the difference between the lowest MP between two consecutive APs (or in the 300ms following the AP, if the ISI interval was too long) and Vrest. Exceptionally, in figure 7 when the AP thresholds between the first and the second AP of the burst were compared, the AHP amplitude was computed relatively to the first AP-prepotential instead of Vrest; this because the threshold of the first AP was more affected by the prepotential than the Vrest (see results). Note that according to our convention AHP amplitudes are negative numbers.

The AHP slope was defined as the slope of a linear regression of MP between the AHP peak and next AP threshold, but restricted to a 20%-60% MP range starting from AHP peak (0%) up to previous AP threshold potential (100%). The MP range for linear regression was indeed based on the preceding AP, in order to allow computing the AHP slope following the last AP of a burst in the same way as the AHP slopes within the burst.

The AHP duration was defined as the period from AHP peak to next AP threshold time. The Intra burst frequency was defined as the average of the inverse ISIs within burst.

We noticed that following a burst, a slow AHP component induced a slow repolarization of the MP toward Vrest. To quantitatively characterize this component for each MC, we selected all bursts with a following interburst interval of at least 500ms. Electrophysiological traces were aligned on the AHP peak of the last burst and a median trace was calculated. We then fitted this median trace from 50ms to 500ms after the AHP peak with a single exponential which gave the slow-AHP time constant. Because of our choice of a minimum interval of 500ms, some cells had no burst selected for median computation, this fit was thus possible in only 42 out of the 49 cells used in this study.

### Threshold model

In order to predict the expected AP threshold following the last burst AP, for each cell, we fitted a model of intra-burst AP threshold as a linear combination of AHP amplitude, AHP slope, AHP duration and Vrest using an Ordinary Least Square regression method. Note that we also tested models taking into account interactions between these parameters but the increase in fit reliability (based on the Bayesian information criterion) was negligible and did not justify taking these interactions into account.

Once a model was fitted for a given cell, we could predict the expected AP threshold following each burst last-AP as a function of time elapsed after the last AHP maximum time (which gave the AHP duration parameter, other model parameters being constant).

### Neuron model

A single compartment model was simulated with NEURON 7.8. All simulations were run with 100-μs time steps. The nominal temperature was 30°C. The voltage dependence of activation and inactivation of Hodgkin-Huxley–based conductance models were taken from (Hu *et al.,* 2009) for Nav and KDR and from (Rubin & Cleland, 2006) for IA. The equilibrium potentials for Na+, K+, and passive channels were set to +90, −91 and −28.878 mV, respectively. We began by constructing a model with the following conductances densities: 0.02 S/cm2, 0.0002 S/cm2, 0.003 S/cm2 and 3.33*10-5 S/cm2 for Nav, KDR, I_A_ and passive channels, respectively. This model a resting membrane potential of −60 mV without holding current injection. In all the other model configurations, we injected a holding current during the simulation to maintain the resting membrane potential at −60 mV.

In Figure 5, no I_A_ conductance was implemented in the model. The conductance density of Nav was set to 0.02 S/cm^2^, 0.005 S/cm^2^ or 0.0025 S/cm^2^. The holding current was set to −30.826 pA, −29.49 pA and −29.26 pA respectively. Spikes were induced by 3ms positive current steps of 400 pA. Hyperpolarizations before APs were induced by 50ms negative current steps whose amplitudes were set in order to obtain a pre-AP membrane potential from −60 mV to −70 mV. AP threshold was defined by the voltage point at which the first time derivative (dV/dt) went above 40 mV/ms. The curve of AP threshold vs. pre-AP activatable Nav conductance was constructed by varying Nav conductance density from 0.04 to 0.0025 S/cm2, inducing one spike from resting membrane potential and measuring AP threshold. The pre-AP activatable Nav conductance values were obtained by multiplying the percentage of non-inactivated Nav conductance - just before the positive current step - by the total Nav conductance density.

In Figure 13, the conductance density of Nav was set to 0.02 S/cm2. The conductance density of I_A_ was set to 0 or 0.003 S/cm2. The conductance of I_A_ was either directly taken from Rubin and Cleland (2006) or modified to get biophysics closer to previously published I_A_ biophysics (Amendola *et al.,* 2012). The modifications were done on the inactivation of I_A_ conductance: modified I_A_ displayed a more depolarized half-inactivation (−90 mV instead of −110 mV), a larger slope of inactivation curve (0.1 mV^−1^ instead of 0.056 mV^−1^) and a shorter inactivation time constant (50ms instead of 150ms). The holding current to keep the resting membrane potential at −60 mV was −30.826 pA for no I_A_ condition, 0 pA for I_A_ condition and −5.85 pA for I_A_ modified condition. Spikes were induced by 3 ms positive current steps of 1 nA. Trains of spikes were induced by trains of these current steps, at 40 Hz.

### Statistical analysis

In many cases, we computed correlations between the different parameters characterizing burst dynamics. We generally computed and plotted the within cell correlations. Summary plots show, for each cell, the slope of the correlation (left part of the figure), the strength of the correlation (R of the linear regression, right part of the figure). Single cell statistically significant correlations (p < 0.05, corrected for multiple comparisons with the Bonferonni-Holm methods) are shown in red (for both the slope and strength of the correlation). Boxes show the mean and 95% confidence interval of the mean.

Global population statistical analysis was performed on the slopes and the correlation coefficient of individual cells using standard t-tests, assessing that the population averages were different from 0. If not stated differently in results, the text gives the mean ± 95% confidence interval (CI, defined as 1.96 x SEM). Other quantities of interest are effect sizes (ES), correlation coefficients (R), t-values (t) and p-values (p) of T-test on R values, and number of cells used in the analyses (N). Note that p-values of T-test performed on slopes are only given in figures. Bayesian analysis was performed with JASP (JASP Team (2020). JASP (Version 0.14.1) [Computer software] by using the default effect size prior (Chaucy scale = 0.707).

### Exclusion criteria

23 spontaneously active MCs where excluded from the analysis based on the following criteria: 5 showing bad quality of recording, 3 showing only tonic activity, 11 showing too few bursts (< 4 bursts), 2 having too high membrane resistance to be identified as MCs (> 500 MΩ), 2 with only too short interburst intervals to compute resting potential (< 200 ms).

### Software

All analyses were done with laboratory custom Python 2.7 scripts, using statistical or curve fitting functions from Scipy 1.2.2, and multiple comparison functions or multiple regression functions from StatsModels 0.9.0

### Data availability

All raw electrophysiological traces, script for traces analysis, data analysis and models scripts are available at Open Science Framwork (https://osf.io/s2dbw/)

## Results

### Heterogeneity of spontaneous firing activity between different MCs

Whole cell recording was performed on 119 MCs in OB slices obtained from 21 rats aged between 30 and 42 days. Firing activity was observed in 72 (60%) MCs at resting MP or after slight membrane depolarization. Among these cells, 23 were excluded from the analysis based on the criteria detailed in the methods. As previously shown (Chen & Shepherd, 1997; Desmaisons *et al.,* 1999; Balu *et al.,* 2004), firing activity was characterized by clusters of APs, henceforth denominated bursts, separated by silent periods presenting subthreshold membrane oscillations (Fig 1A). A total of 1532 bursts (with at least 2 APs) and 386 isolated APs were analysed. Burst properties such as the number of APs, membrane potential at which bursts occurred, inter-burst frequency and intra-burst frequency were heterogeneous (Fig 1B, 1C, 1D and 1E; left). Such a heterogeneity could be partly due to specific differences among the recorded MCs (Fig 1B, 1C, 1D and 1E; right) which population shows intrinsic biophysical diversity (Padmanabhan & Urban, 2010). But it was also partly due to the difference of the average holding potential (Vrest) between the different MCs. In fact, more depolarized MCs presented higher intra-burst frequency (R = 0.58, Wald test, p < 0.001, N = 49) and larger burst size (R = 0.38, Wald test, p=0.007, N=49). Burst size and intra-burst frequency (R = 0.56, Wald test, p < 0.001, N = 49) were also positively correlated.

**Fig 1:**
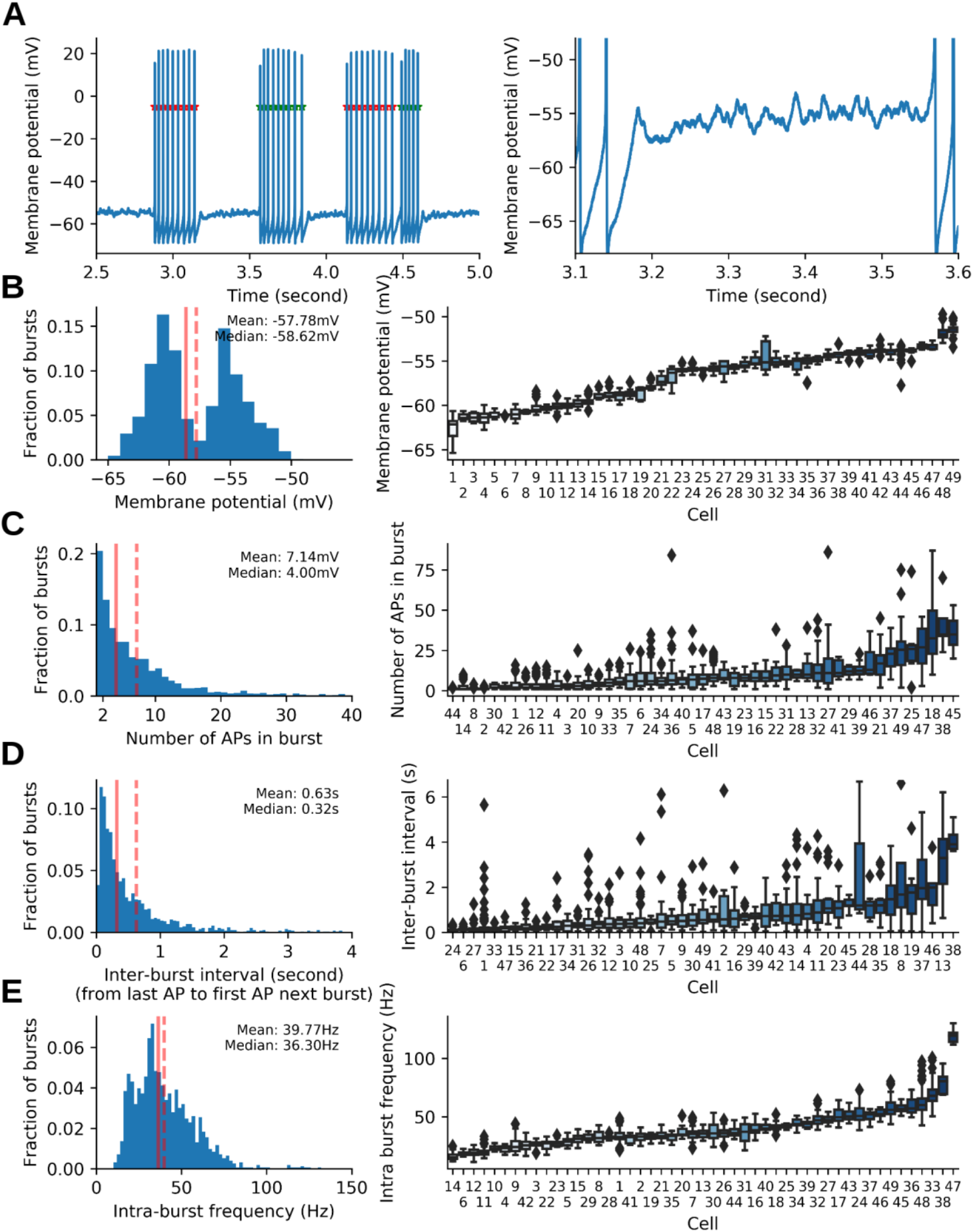
Burst properties of MCs. **A:** Left, example of spontaneous recording in MCs showing typical bursts of APs. APs belonging to the same burst are alternatively marked with red and green stars. Right, enlargement of the trace in A left which shows MP dynamics with fast oscillations which were typical during interburst periods. **B:** Distribution of MP values preceding bursts (Vrest, **left:** for all detected bursts, **right:** whisker plot per cell showing inter-cell variability); the distribution is bimodal because some cells were spontaneously active while others not and needed to be slightly depolarized to spike. **C:** Distribution of burst sizes (same representation as in B). **D:** Distribution of inter-burst intervals (same representation as in B). **E:** Distribution of intra-burst frequency (same representation as in B). Continuous and dashed vertical lines in left panels materialize the mean and median values respectively.

### Dynamic modulation of AP threshold at resting potential

It has been reported that, in MCs, the AP threshold decreases when firing is preceded by a MP hyperpolarization induced by current injection (Balu *et al.,* 2004). Interesting is that, spontaneously, burst initiation was ever preceded by a MP hyperpolarization (see an example in Fig 2A) being in agreement with a previous report (Desmaisons *et al.,* 1999). We therefore investigated whether oscillatory activity, potentially associated with spontaneous inhibitory transmission, was capable to produce a dynamic modification of AP threshold, thus to contribute to the firing initiation. In cortical neurons, the threshold is affected by the trajectory of MP that precedes the AP. In particular, when the MP was hyperpolarized or the rate of membrane depolarization (dVm/dt) preceding the AP was faster, more negative AP threshold were observed (Henze & Buzsáki, 2001; Azouz & Gray, 2003; Li *et al.,* 2014). Similarly, in MCs, the first AP threshold of a burst positively correlates with the value of MP calculated at the negative peak of the oscillation preceding AP (thereafter named AP-prepotential, see example in Figure 2B, and population analysis in Figure 2C). The average modification of AP threshold was of −0.37 ± 0.94 mV for 1 mV of MP hyperpolarization (ES = 1.13, R = 0.48 ± 0.08, T-test t = 11.6, p < 0.001, N =49), meaning that AP threshold was driven towards more negative values by membrane hyperpolarization. However, contrary to what was reported for cortical neurons and predicted by theoretical models (Platkiewicz & Brette, 2011), in MCs, the AP threshold was less negative when the membrane-depolarization rate (dVm/dt) preceding AP was fast, according to average modification of AP threshold of 0.85 ± 0.71 mV per 1 mV/ms of membrane depolarization (Fig 2D, ES = 0.34, R = 0.18 ± 0.07, T-test t = 4.96, p < 0.0001, N = 49). This unexpected effect may be a consequence of the small positive covariation we measured between the AP-prepotential and the pre-AP slope (lower pre-AP slope for more hyperpolarized AP-prepotential; average slope 0.04 ± 0.02 (mV/ms)/mV, ES = 0.52, R = 0.13 ± 0.07, T-test t = 3.41, p = 0.0013, N = 49, not shown). According to the literature, this covariation should induce opposite effects on the threshold potential. Here, the prepotential effect appeared to prevail on the pre-AP slope effect. Since the AP threshold of MCs can dynamically shift depending on the recent history of MP, it is conceivable that the firing could be induced by hyperpolarizing events bringing the AP threshold below the median resting potential (Vrest, see Methods) or within the range of subthreshold MP oscillations. This was indeed the case for 27% of recorded bursts (representing 92 % of recorded MCs). Unsurprisingly the level of hyperpolarization (AP prepotential - Vrest) preceding the burst was higher in these cases (see example in Fig 2E left) than in bursts where the threshold of the first AP remained above the Vrest (see example in Fig 2E right): −1.30 ± 0.17 mV vs −0.56 ± 0.07 mV, T-test: t = −9.48, p < 0.001, ES = 0.44, N = 513 and 1405. Altogether these data suggest that spontaneous firing in MCs could be triggered according to two modalities: 1- a classical membrane depolarization above the Vrest, eventually produced by the excitatory synaptic activity; 2- a membrane hyperpolarization produced by the oscillatory and/or inhibitory synaptic activity that would bring the AP threshold below Vrest or within Vrest variability.

**Fig 2:**
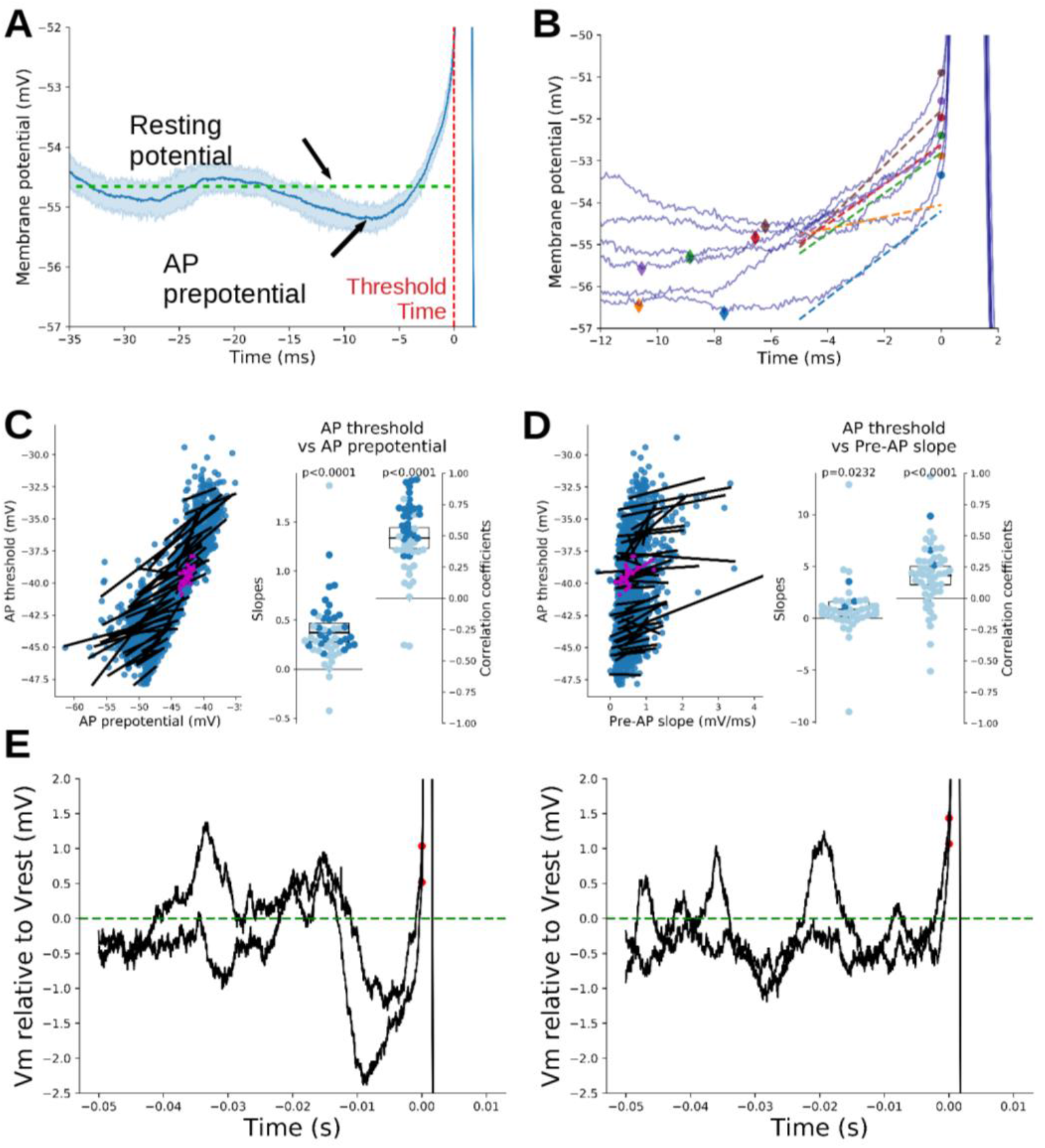
The threshold of the first AP of the burst is affected by the trajectory of the preceding MP. **A:** Burst were initiated after an hyperpolarization phase of the MP. Here, MP traces preceding each burst of a single neuron (#32) were averaged (blue line) along with their STD (shaded area). **B:** Same cell example of the variability of AP prepotential and its correlation with first AP threshold. Each colour spot marks the most negative values of pre-potential (AP pre-potential, diamond) and the respective AP threshold (round dot). Lines of the same colour show the pre-AP slopes of the MP (lines are displayed in the time interval where MP was fitted). See methods for the details of each measure on MP traces. **C:** AP threshold positively correlated with AP pre-potential. Left panel, correlations between the AP-prepotential and first AP threshold within cell bursts. A linear fit was done for each neuron (one dot per burst, the fit is shown as a black line). In purple the measurements and fit of the neuron exemplified in panels A-B. Right panels: scatter plots of the slopes (left) and correlation coefficients (right) obtained from the linear fit done for each neuron in the left panel. Darker dots correspond to individual fits with p < 0.05 (Pearson correlation, corrected for multiple comparisons). Black boxes show the mean and its 95% confidence interval for each distribution (population level analysis). The significance of deviation from 0 of each distribution was further tested with a one sample t-test (p-values are displayed above each scatter plot, see main text and methods for additional details). **D:** Small positive correlation between the MP slope preceding the first AP in bursts (pre-AP slope) and its threshold. Data are presented as in C. **E:** Left, examples of strong membrane hyperpolarization bringing the AP threshold within the range of membrane fluctuations. Right, examples of AP threshold that remained above membrane fluctuations. For a better comparison, traces were aligned on their resting MP (set at 0 for the figure).

### Cellular mechanisms of AP threshold modification

The relationship between MP and AP threshold was further investigated through the experiment depicted in figure 3. Here MCs were slightly hyperpolarized with a steady current injection to prevent spontaneous firing and 2 successive APs were evoked at 6 second intervals by short (3 ms) depolarizing current steps; the second AP being preceded by 50 ms hyperpolarizing current step (Fig 3A, left). The comparison of AP thresholds between the two evoked APs showed that pre-AP membrane hyperpolarization produced a linear shift of AP threshold toward more hyperpolarized values (−0.32 ± 0.05 mV threshold shift per each mV of membrane hyperpolarization; Fig 3A, right, N = 75). The same effect was obtained with pre AP-hyperpolarization durations varying over a range from 10 to 90 ms (Fig 3B, N = 16). The contribution of the membrane depolarization rate to AP threshold was evaluated by comparing the effect of depolarizing ramps at different dVm/dt (Fig 3C, left). For the 11 recorded MCs, the higher the depolarization rate the more negative the AP thresholds (Fig 3C, right). The slope of the linear regression obtained from AP threshold/ depolarizing speed analysis was used to quantify the effect of membrane speed on threshold for each MC (Fig. 3C, inset). This analysis showed a shift of AP threshold of −7 ± 3 μV for each mV/s of modification of membrane depolarizing speed (Fig 3 inset; ES = −1.2 p = 0.003). Therefore, similarly to what was observed in cortical neurons (Henze & Buzsáki, 2001; Azouz & Gray, 2003; Li *et al.,* 2014), the AP threshold in MC became less negative when MP was depolarized with a slow depolarization rate. This result supports the interpretation that the opposite correlation between AP threshold and pre-AP slope (shown in fig 2D) would be a consequence of a more hyperpolarized AP pre-potential when the pre-AP slope is lower.

**Fig 3:**
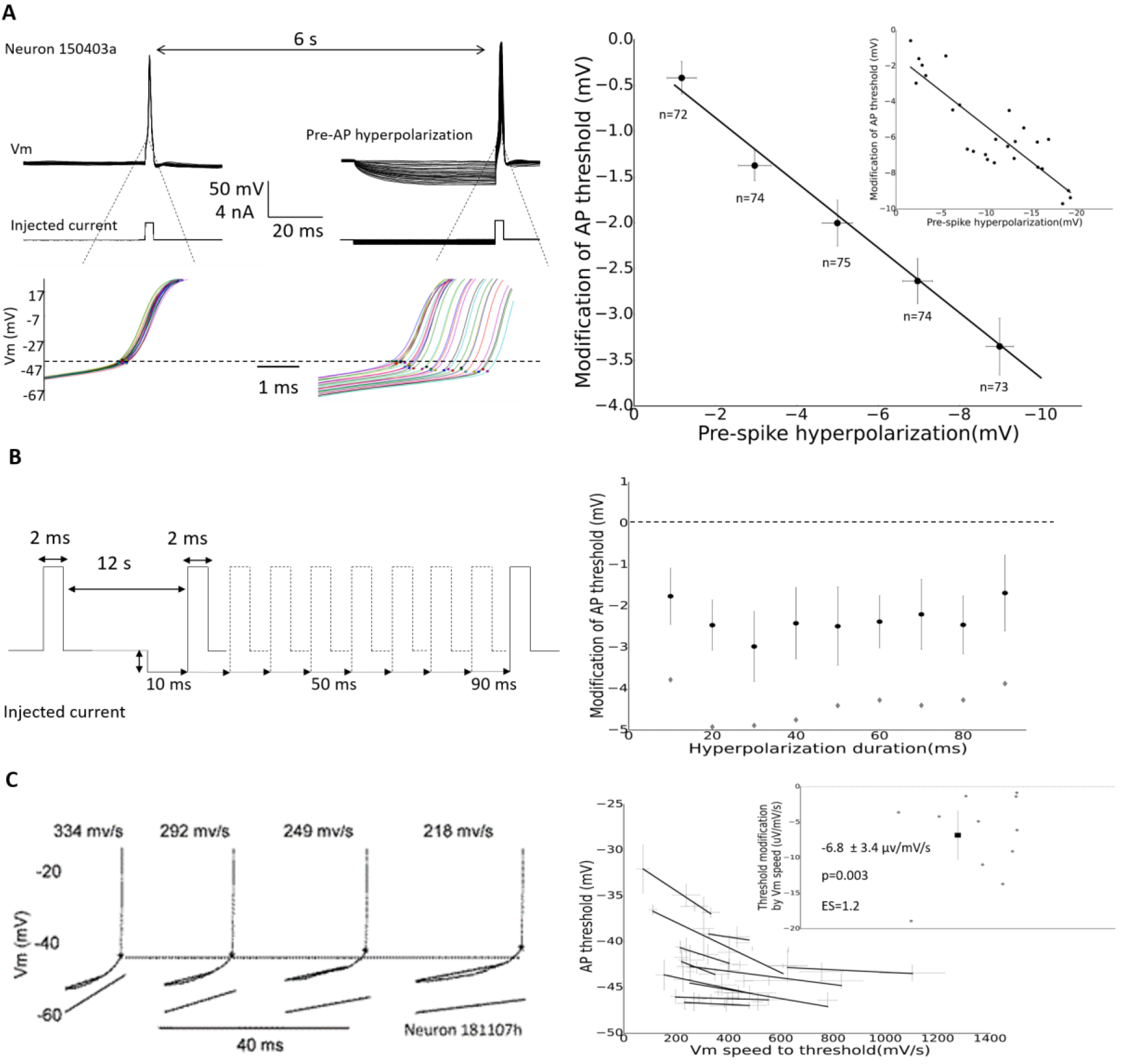
AP threshold of MCs decreased with membrane hyperpolarization and high depolarization speed. **A)** AP threshold decreased proportionally with membrane hyperpolarization. Left; experimental protocol and representative example of threshold modification by membrane hyperpolarization. Dots set threshold position, dashed line sets the threshold position in absence of hyperpolarization. Right; quantification of spike threshold modifications produced by membrane hyperpolarization. Inset; analysis performed on the neuron 150403a depicted on the left. Number of cells is in brackets. **B)** Spike threshold modification was not affected by the duration of membrane hyperpolarization. Left; schematic representation of the experimental protocol. Right; quantification of the effect. Black circles: average AP threshold modification; grey diamonds: average membrane hyperpolarization (same scale than threshold modification). **C)** AP threshold decreased with the membrane depolarization rate. Left; experimental protocol and representative examples of AP thresholds for different depolarization rates. Asterisks set the threshold position; lower lines materialize depolarization current ramps (only the last 10 ms preceding the threshold is shown). Right; quantification of AP threshold as a function of membrane depolarization rate for 11 MCs. Inset quantification of AP threshold modification as a function of membrane depolarization rate for the 11 MCs. ES = effect size. Horizontal and vertical bars represent 95% Confidence Interval.

The most likely mechanism responsible for the modification of threshold produced by membrane hyperpolarization is the recovery from inactivation of voltage dependent channels implicated in AP generation; namely, the Na^+^ and T-type Ca^2+^ channels. To test this assumption, we used the same experimental procedure as described in figure 3A, combined with a pharmacological approach (figure 4A). Since the pharmacological compounds were applied at different time periods, we first made sure that the effect of membrane hyperpolarization on AP threshold effect was stable over time. As shown in figure 4B a time dependent spontaneous shift of AP threshold toward more depolarized values was observed during whole cell recording in control condition, possibly due to cell dialysis (see Ctr 5 min and Ctr 10 min). Despite this, the effect of membrane hyperpolarization on the modification of the AP threshold was not affected (Fig 4 C,D). The participation of T-type Ca^2+^ channel was investigated by using the selective antagonist ML218 (5-10 μM). Although, the shift of AP threshold compared to Ctr condition was slightly more important than that produced by cell dialysis at 5 min it did not reach significance (Fig 4B; 4.1 ± 2.4 mV; ES = 0.5, p = 0.16 compared to Ctr at 5 min).

**Fig 4:**
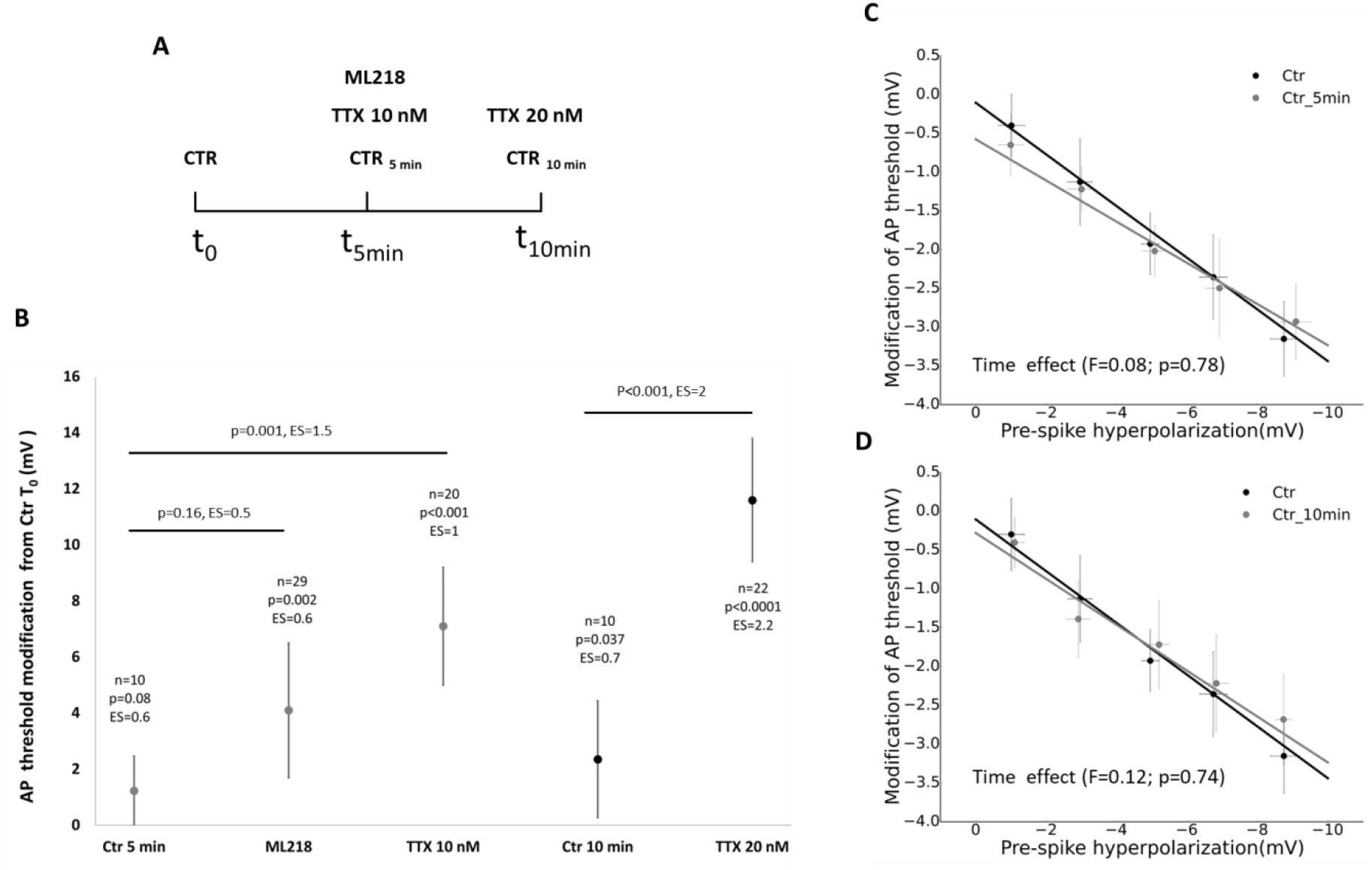
Modification of MC AP threshold and hyperpolarization effects over time and after applications of Na^+^ and Ca^2+^ channels antagonists. **A)** Timing of experiments during which AP threshold was measured after 5 and 10 min, in control conditions or after different pharmacological applications. **B)** AP threshold was significantly increased by blockade of Na^+^ channels with TTX, but not by the antagonist of T types Ca ^2+^ channels (ML218: 5-10 μM). Note the spontaneous temporal increase of threshold in control condition, both at 5min and 10 min, with a small difference, possibly due to cell dialysis. The statistical analysis compared to Ctrt0 is depicted above the data points. **C, D)** The effect of membrane hyperpolarization on the AP threshold was stable with time. ES = effect size. Horizontal and vertical bars represent 95% Confidence Interval.

As show in figure 5A ML218 did not modify the shift of AP threshold induced by membrane hyperpolarization (Fig 5A; threshold/hyperpolarization slope modification −0.0 ± 0.1 mV/mV, ES = 0.1, p = 0.66; N = 21) indicating that this effect was not based on recovery from inactivation of T-type Ca^2+^ channels. To investigate the effect of Nav channels recovery from inactivation on AP threshold, we first used a toy model containing only one Nav channel type and one Kv delay-rectifier channel type. This model reproduced the shift of AP threshold produced by the membrane potential hyperpolarization (Fig 5B-C, 200 pS/m^2^). Such an effect was attributable to the recovery from inactivation of Nav channels produced by the MP hyperpolarization, leading to a hyperpolarization of the AP threshold. Indeed, as expected, decreasing the density of Nav channels in the model, leaded to a shift of AP threshold toward more positive values (Fig 5B). More interesting, under reduced Nav condition, the hyperpolarization of membrane potential had a bigger effect on AP threshold than in control condition (Fig 5C). This can be easily explained by taking into account that AP threshold is not linearly correlated with the quantity of activatable Nav channels. Fig 5D shows the curve of the AP threshold vs the quantity of activatable Nav conductance just before the AP: here the quantity of Nav conductance activatable at resting MP (−60 mV) or after 10 mV hyperpolarization (−70 mV) was plotted for the 3 conditions of Nav channels density (200 pS/m^2^ in red, 50 pS/m^2^ in green or 25 pS/m^2^ in blue). We can see that the increase of pre-spike Nav channels availability produced by MP hyperpolarization produced a greater shift in AP threshold in reduced Nav condition (50 pS/m^2^ or 25 pS/m^2^) than in control condition (200 pS/m^2^). The prediction of the model was checked by applying low doses of the Na channel blocker, TTX. As shown in figure 4B TTX at 10 nM shifted the AP threshold to more positive value (+7.1 ± 2.1 mV, ES = 1, p < 0.001, compared to Ctr 5 min) while a similar more pronounced effect was observed at 20 nM (+11.6 ± 2.2 mV, ES = 2, p < 0.001, compared to Ctr 10 min). These observations support that AP threshold depended well on the density of Na^+^ channels potentially de-inactivable by hyperpolarization. As predicted by the model, with TTX at 20 nM the effect of membrane hyperpolarization on AP threshold was amplified (Fig 5E; increase of threshold/hyperpolarization slope 0.26 ± 0.18 mV/mV, ES = 0.6, p = 0.01, N = 22). Interestingly, this effect was not observed with TTX at 10 nM (Fig 5F; threshold/hyperpolarization slope modification −0.03 ± 0.18 mV/mV, ES = −0.16, p = 0.48, BF_10_ = 0.3, ‘evidence of absence’, N = 19), despite the fact that reduction of Nav channels availability appeared sufficiently important, as shown in fig 4B. This result suggests that, around −60 mV, the recovery from inactivation produced by the hyperpolarization would mainly involve the Nav channel sub-types having a lower affinity for the TTX (see discussion).

**Fig 5:**
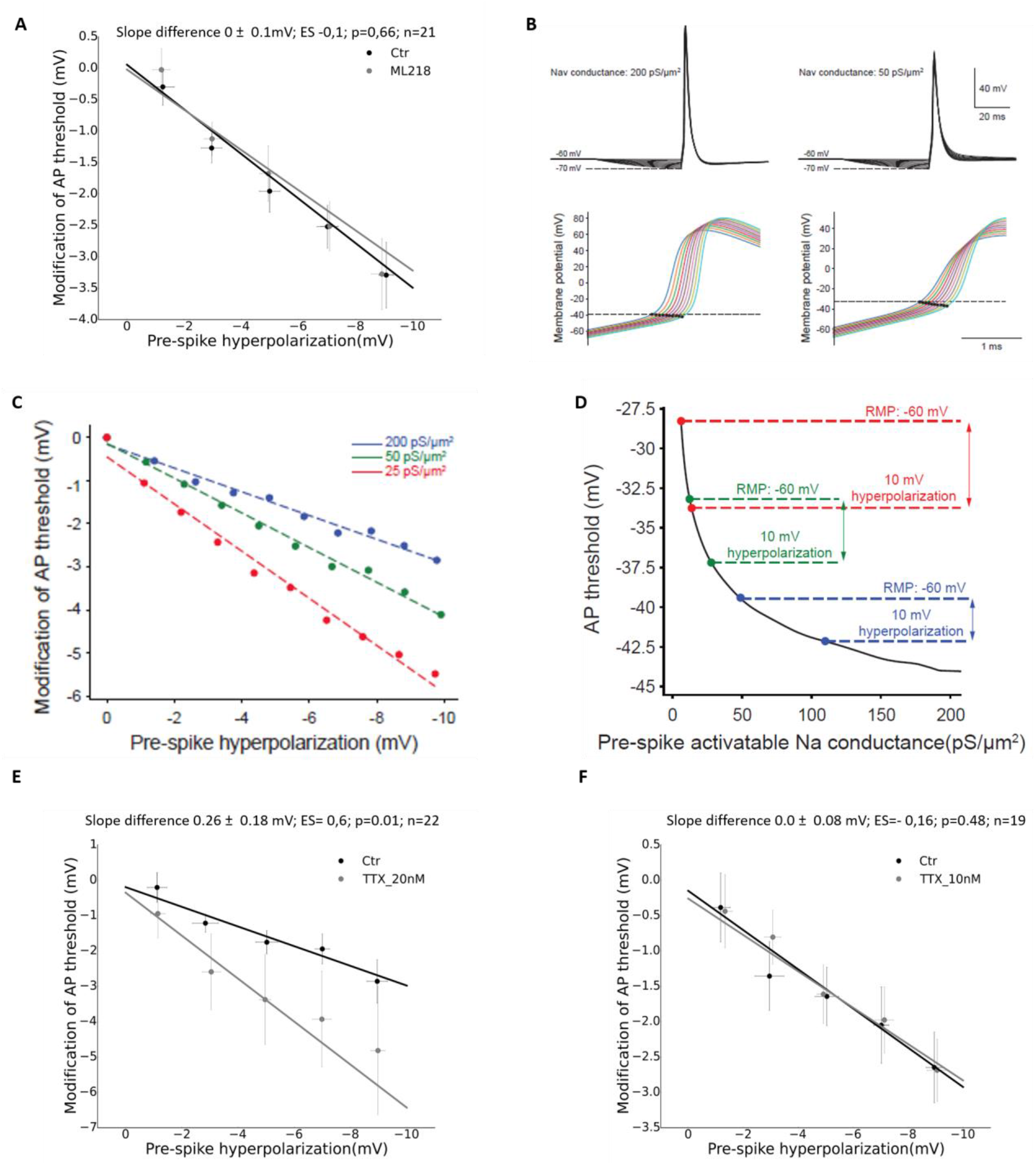
Sodium but no calcium channel antagonists amplified the effect of membrane hyperpolarization on AP threshold. **A)** The shift of AP threshold produced by membrane hyperpolarization was not modified by the antagonist of T-type calcium channel ML218 (5-10 μM). **B)** Toy model simulation showing the modification of AP threshold with membrane hyperpolarization and availability of Nav channels. **C)** Quantification of the effect of membrane hyperpolarization and Nav channels availability on AP threshold, showing an amplification of the AP threshold shift induced by membrane hyperpolarization when the density of Nav channels was reduced. **D)** Curve depicting the modification of AP threshold as a function of the quantity of Nav conductance available before the AP. Note that the same 10 mV membrane hyperpolarization produced a stronger modification of AP threshold when the density of Nav channels was reduced (compare the 3 conditions: 200 pS/m^2^, 50 pS/m^2^, 25 pS/m^2^) **E)** TTX at 20 nM amplified the AP threshold shift induced by membrane hyperpolarization. **F)** TTX at 10 nM failed to amplified the AP threshold shift produced by membrane hyperpolarization ES = effect size. Horizontal and vertical bars represent 95% CI.

We finally determined whether, in MCs recorded close to resting MP (−60 mV), short hyperpolarizations of Vm could produce a recovery from inactivation of Na^+^ channels. To this end, we performed the experiment depicted in figure 6. Here, Na^+^ current was pharmacologically isolated (see Methods) and MCs were recorded in voltage clamp configuration. Following the short membrane pre hyperpolarization (25 ms), the amplitude of Na^+^ current generated by a 5 ms depolarization step to −10 mV, increased, proportionally to the hyperpolarization level (Fig 6B, p=0.0002, N=6, Friedman test). Moreover, the recovery from inactivation of Na^+^ channels was independent of the duration of the hyperpolarization, in the 10 to 90 ms range, (Fig 6C, p = 0.74, N=6, Friedman test). These results echoed the effects produced by membrane hyperpolarization on AP threshold that were depicted in figure 3A and 3B, further supporting the hypothesis that the threshold shift was based on the recovery from inactivation of Na^+^ channels.

**Fig 6:**
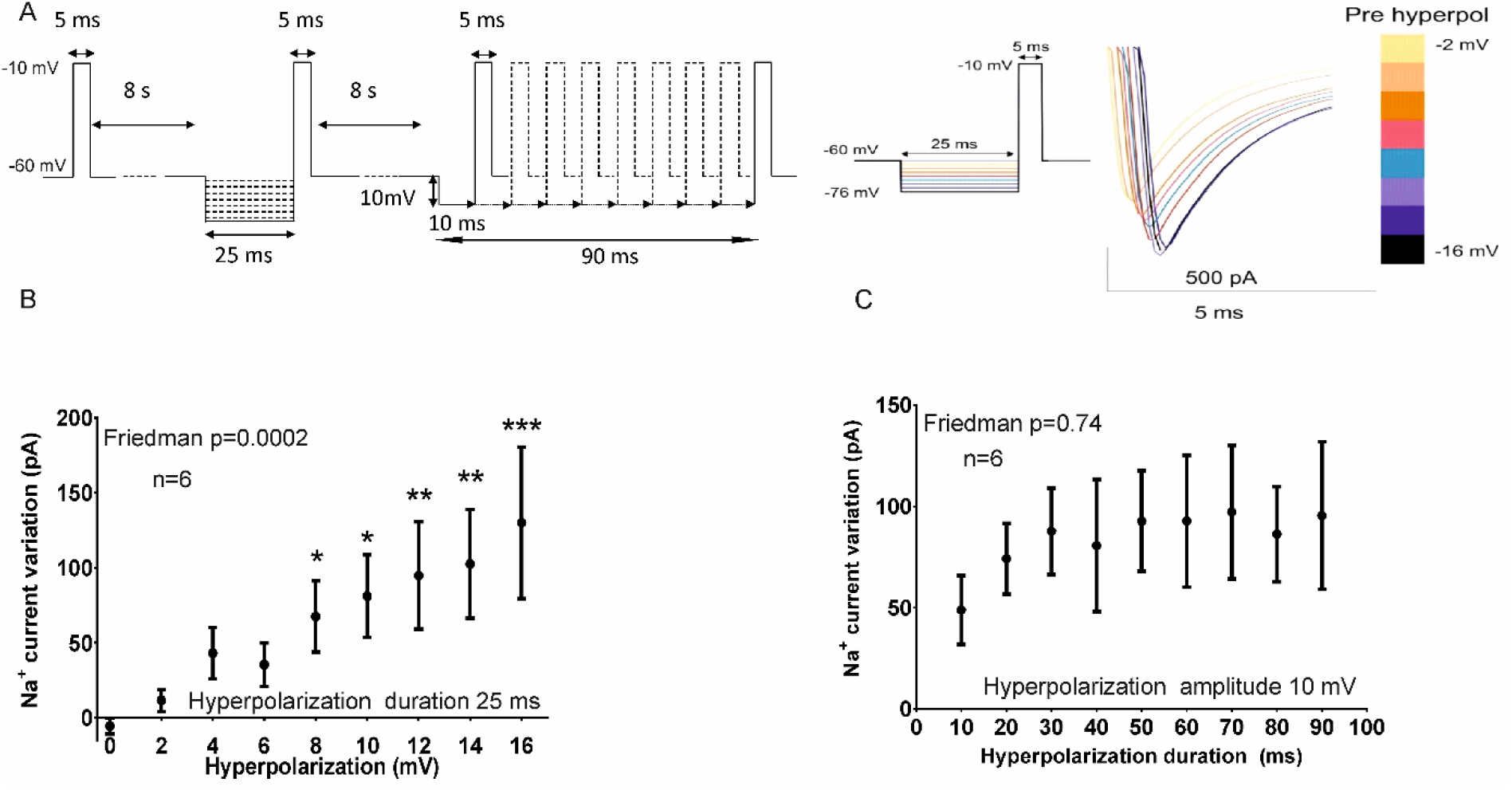
Membrane pre hyperpolarization resulted in an amplification of Na^+^ current in MCs. **A)** Left; experimental protocol depicting the imposed modifications of membrane voltage. Right, representative of Na^+^ current following membrane hyperpolarization. **B)** Quantification of the modification induced by different levels of membrane hyperpolarization on Na^+^ current amplitude. **C)** Quantification of the modification produced, by different duration of membrane hyperpolarization, on Na^+^ current amplitude. * p<0.02; ** p<0.01; *** p<0.001; post hoc comparisons with the condition without hyperpolarization, Dunn’s test. Vertical bars = sem.

### Spontaneous bursting activity was promoted by dynamic modification of AP threshold induced by the AHP

We next investigated the impact of dynamic modifications of AP threshold on MC firing.

As shown in Figure 7 A-B, the AP thresholds within the bursts were naturally shifted to more negative potentials than those of the first AP (delta threshold between the first and second AP = −2.28 ± 0.09 mV, N = 1532 bursts, ES= −1.28, T-test t = −50.0, p < 0.001). The most likely explanation of the threshold shift within the burst is that it was due to membrane hyperpolarization induced by the preceding AHP. Indeed the modification of AP threshold, calculated between the first and the second AP of the bursts, positively correlated with the AHP amplitude measured relatively to the first AP prepotential (−0.35 ± 0.10 mV threshold shift for each mV of membrane hyperpolarization, ES = −1.03, R = −0.42 ± 0.09, T-test t = 9.34, p < 0.0001, N = 49, Fig 7C) as well as, with the repolarization rate of the AHP (AHP slope) (−5.6 ± 1.5 mV for each mV/ms of modification of MP rate, R = −0.47 ± 0.09, ES = −1.04, T-test t = −10.4, p < 0.0001, N = 49, Fig 7D). These results suggested that both the AHP amplitude and repolarization rate accounted for the negative AP threshold shift within the burst. Interestingly this effect, observed at the single cell level, was also seen at the population level looking at average cell values (average delta AP threshold vs average MP hyperpolarization: slope = −0.33 mV/mV, R = −0.57, Wald test, p < 0.001, N = 49; average delta AP threshold vs average modification of MP repolarization rate: slope −3.4 mV/(mV/ms), R = −0.7, Wald test, p < 0.001, N = 49; data not shown).

**Fig 7.**
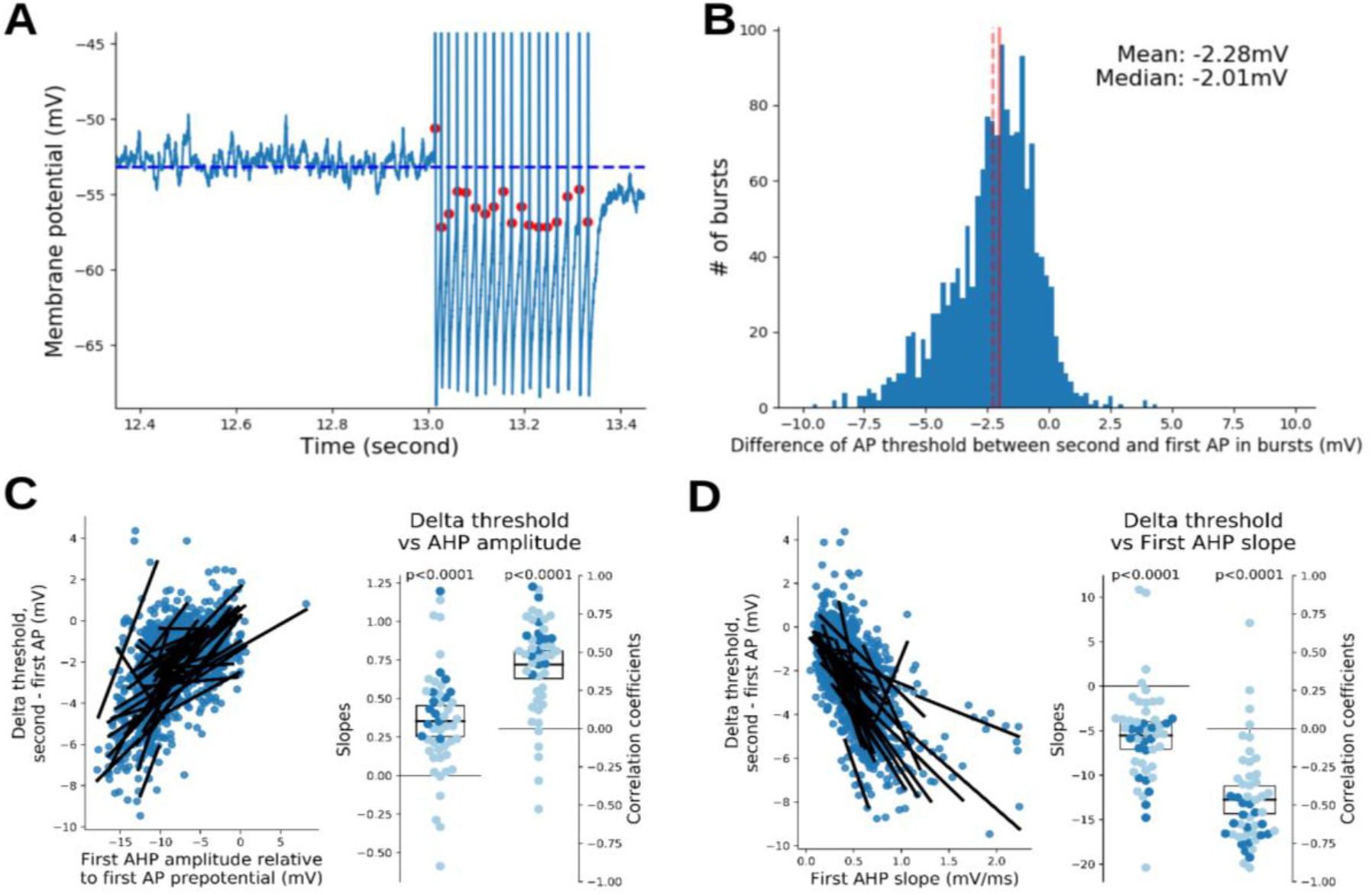
The AHP induced hyperpolarization lowered the AP threshold within the bursts. **A:** Left, example illustrating the lowering of threshold for APs within the burst. **B:** Histogram of differences of AP threshold potentials between the second and the first AP of the bursts. **C:** Larger AHP produced higher hyperpolarization of AP threshold. Positive correlation between the difference of the second and first AP threshold and the first AHP amplitude (computed here relatively to the first AP prepotential). Results are presented as in Figure 2C. **D:** Faster AHP repolarization produced higher hyperpolarization of AP threshold. Negative correlation between the difference of the second and first AP threshold and the first AHP slope. Results are presented as in Figure 2C.

Once the first AP fired from resting potential, AP threshold shift induced by AHP could potentially act as regenerative mechanisms of firing, especially in the case where it would drive the threshold below the resting MP (Fig 7A) or within the range of spontaneous membrane oscillations (Fig 8A). To precisely assess the position of AP thresholds relatively to subthreshold fluctuations, we normalized the AP thresholds inside the bursts (i.e. all the APs except the first one) using a linear interpolation. We set the Vrest at 0 to and the maximum amplitude of the subthreshold fluctuations, at 1. In such conditions, a negative value for the normalized AP threshold meant that it was more hyperpolarized than Vrest, a value between 0 and 1 meant that it was within the range of MP intrinsic subthreshold fluctuations and a value above 1 meant that it was more depolarized than subthreshold fluctuations. As shown in figure 8C, AP threshold was more negative than Vrest in 39 % of bursts: it remained within the range of subthreshold membrane oscillations for 52 % and above membrane oscillatory activity for the remaining 9 % (Fig 8C). A clear heterogeneity of the AP threshold value was observed between MCs, with respect to Vrest (Fig 8D). Interestingly the AP threshold shift relative to the Vrest (relative AP threshold), appeared to affect bursts properties. Stronger negative shifts were associated to burst having higher number of APs (correlation between burst size and relative AP threshold: average slope = −4.87 ± 2.22 AP/mV, ES = −0.62, average R = −0.37 ± 0.08, T-test t = −9.56, p < 0.0001, N = 49, Fig 8E1) and higher intra-burst firing frequencies (correlation between bursts frequency and relative AP threshold: average slope = −4.46 ± 1.00 Hz/mV, ES = −1.26, average R = −0.59 ± 0.10, T-test t = −11.9, p < 0.0001, N = 49 Fig 8F1). Similar correlation was observed between the average AP threshold modification in different MCs and their average bursts size and intra-burst frequency (see Fig 8E2 and 8F2), suggesting that the heterogeneity of MC firing properties shown in figure 1 could be, at least partly, ascribable to AP threshold modification induced by AHP.

**Fig 8:**
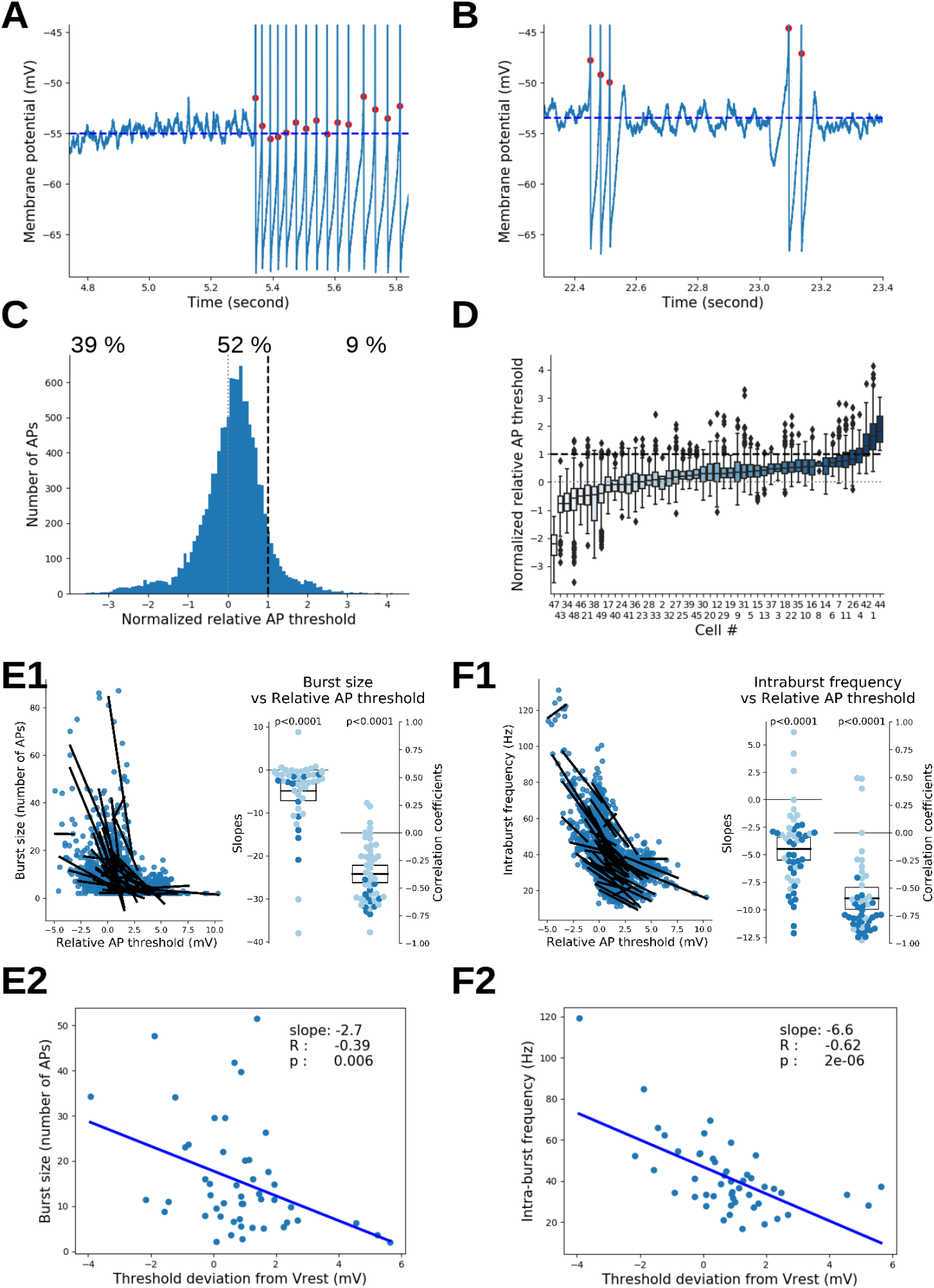
The shift of AP threshold determined the firing properties of the bursts. **A:** example of intraburst AP thresholds that remained within the range of MP oscillations. **B:** example of intraburst AP thresholds shifting above MP oscillations. Note that the rebound of MP after the AHP could potentially allow the burst generation despite high AP threshold relative to Vrest **C:** distribution of the AP threshold relative to Vrest (first burst APs excluded) normalized so that 1 (dashed line) corresponds to the maximum MP reached during rest. **D:** same analysis as in **C** for each MC. **E:** *higher* hyperpolarization of *AP threshold was associated with bursts having higher number of APs*. **F**: *higher* hyperpolarization *of AP threshold was associated with bursts* having higher mean intraburst frequencies. **E1** and **F1** are within cell analysis, results being presented as in Figure 2C. **E2** and **F2** are between cell analysis (lines: linear fits of average burst size or average burst frequency vs average relative AP threshold, R: correlation coefficient, p: Wald -test p-value for the slope, N = 49)

As expected from the relationship between AHP and AP threshold shift, the intra-burst firing properties correlated with AHP properties. Higher AHP amplitude were associated to longer bursts (correlation between burst size and AHP amplitude: average slope=-2.98 ± 1.92 AP/mV, ES = −0.43, average R = −0.13 ± 0.12, T-test t = −2.26, p = 0.03, N = 49, Fig 9A1) and higher firing frequencies (correlation between firing frequencies AHP amplitude: average slope −3.31 ± 1.36 Hz/mV, ES = −0.69, average R = −0.33 ± 0.12, T-test t = −5.23, p < 0.0001, N = 49; Fig 9B1). In addition, faster AHP repolarizations were associated with longer bursts (correlation between burst size and AHP slope: average slope = 38.3 ± 22.7 AP/(mV/ms), ES = 0.48, p = 0.002; average R = 0.24 ± 0.08, T-test t = 5.79, p < 0.0001, N = 49, Fig 9C1) and higher firing frequencies (correlation between intra-burst frequency vs AHP slope: average slope= 71.0 ± 9.0 Hz/(mV/ms), ES = 2.24; average R = 0.76 ± 0.07, T-test t = 21.7, p < 0.0001, N = 49, Fig 9D1). Therefore, the speed of repolarization of the AHP appeared to more strongly impact MC firing properties than the AHP amplitude. Similar contribution of the AHP to the MC firing properties were observed when performing between cell analyses (see Fig 9, panels A2, B2, C2, D2); suggesting that firing heterogeneity between the different MCs reported in figure 1 are, at least partly, based on the heterogeneity of AHP characteristics.

**Fig 9:**
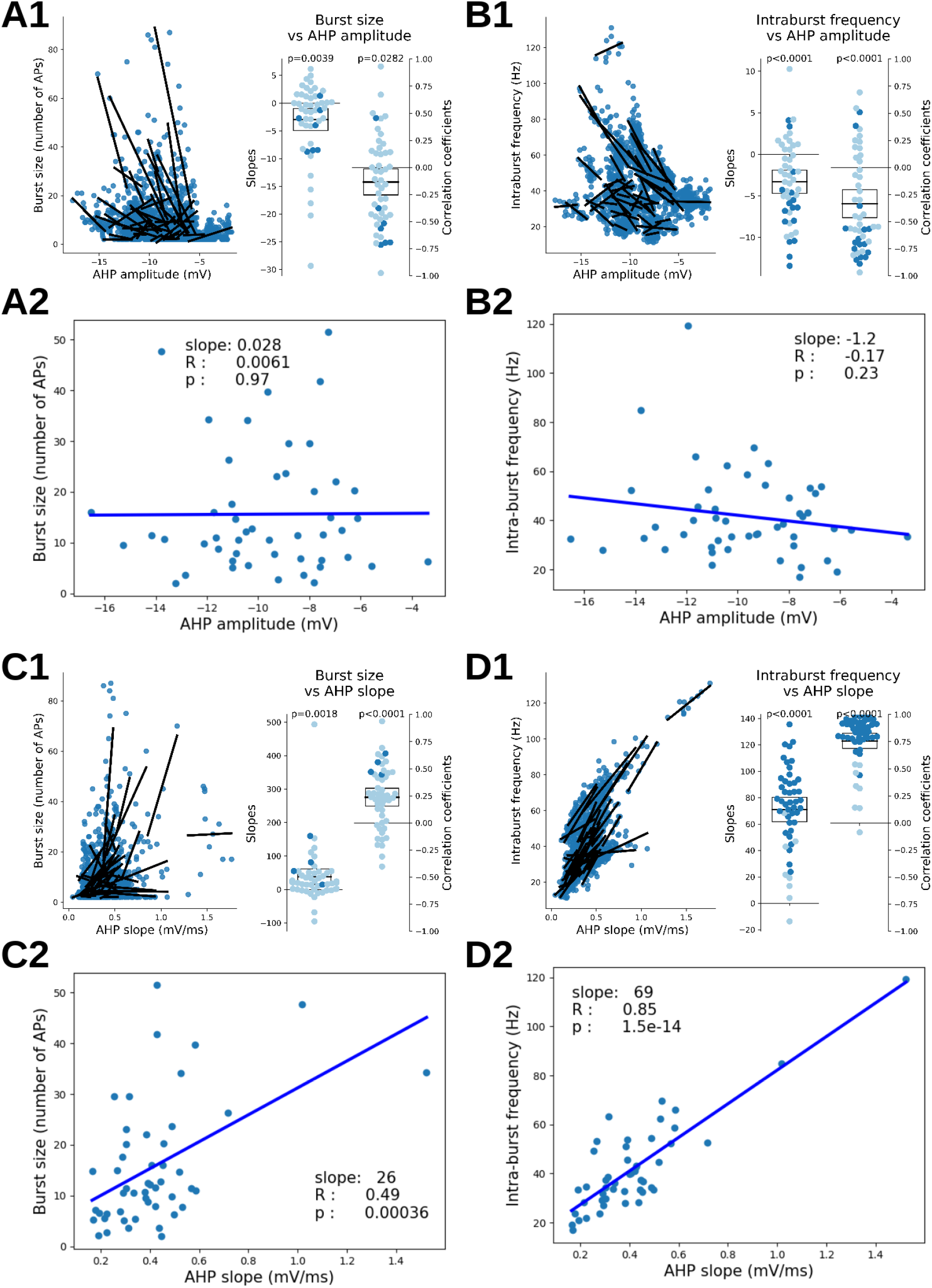
AHP characteristics determined the firing properties of bursts. **A:** Larger AHP amplitudes were associated with longer bursts. **B:** Larger AHP amplitudes were associated with higher intraburst frequency. **C:** Faster AHP repolarizations **were** associated with longer bursts. **D:** Faster AHP repolarizations **were** associated with higher intraburst frequency. Note that correlation coefficients were higher when the AHP slope was considered with respect to the AHP amplitude. In panels **A1**, **B1**, **C1** and **D1** (within cell analyses), results being presented as in Figure 2C. In panels **A2**, **B2**, **C2** and **D2** (between cell analyses); results being presented as in Figures 8E2 and 8F2.

So far, we provided evidence that the frequency and duration of MC bursts were affected by the negative shift of AP threshold produced by the AHP, through the recovery from inactivation of Na+ channels. These two parameters seemed also MP dependent since both increased with membrane depolarization (correlation between burst size and MP: average slopes =3.95 ± 4.01 AP/mV, ES = 0.28, average R = 0.32 ± 0.10, T-test t = 6.06, p < 0.0001, N = 49; Fig 10A; correlation between intra-burst frequency and MP: average slopes =2.90 ± 2.41 Hz/mV, ES = 0.34, average R = 0.37 ± 0.13, T-test t = 5.49, p < 0.0001, N = 49, Fig 10B). This effect was likely due to the stronger negative shift of AP threshold between the first and the second AP of the burst when the MP was depolarized (correlation between relative AP threshold and MP: average slopes= −0.97 ± 0.29 mV/mV, ES = −0.94, average R = −0.64 ± 0.10, T-test t = −12.3, p < 0.0001, N = 49, Fig 10C). Three factors could support the latter effect: 1 - an increase of AHP amplitude with MP depolarization (correlation between AHP amplitude vs MP: average slopes= −0.59 ± 0.55 Hz/mV, ES = −0.30, average R = −0.51 ± 0.14^’ T-test t = −7.32, p < 0.0001, N = 49, Fig 10D^), possibly due the increase of K+ driving force with the membrane depolarization; 2 - an increase of the repolarization speed of the AHP with MP depolarization (correlation between AHP slope and MP: average slopes= 0.036 ± 0.014 (mV/ms)/mV, ES = 0.73, average R = 0.33 ± 0.11, T-test t = 5.75, p < 0.0001, N = 49, figure 10E); 3 – a stronger effect of Nav channels recovery from inactivation, due to steady-state-inactivation increase when the MP was depolarized. Indeed, as shown by the toy model (Fig 10 G,H), reducing the quantity of pre-spike activatable Nav channels led to a greater effect of MP hyperpolarization on AP threshold shift. The third factor is experimentally supported by the observation that the slope of the correlation between relative AP threshold and the AHP amplitude, was more important when MCs were depolarized (i.e., for a same AHP amplitude, AP threshold modification increased with MP depolarization) (Fig 10F). Overall, the three factors could cooperate to make the relative AP threshold more negative, thus making the bursts longer and faster for more depolarized MP. All these effects, shown here at the cell level were also seen at the population level (between cell analysis, not shown).

**Fig 10:**
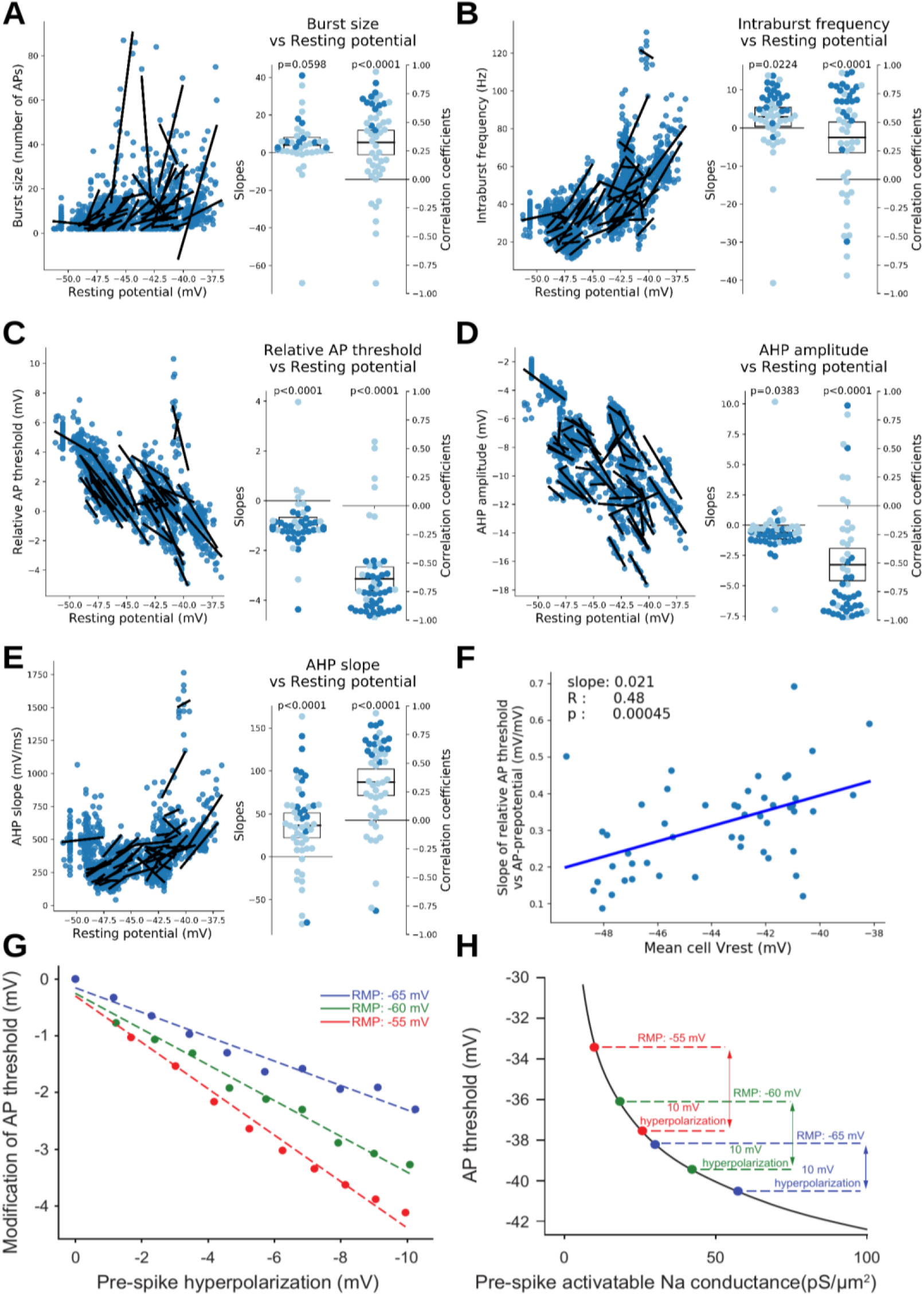
Influence of resting potential on burst size and frequency was linked to the modification of AP threshold, through changes in AHP characteristics and in sodium channels inactivation rates. **A-E:** Linear fits, slopes and coefficients of the correlations between burst firing properties or burst AHP characteristics, with the resting potential preceding the burst; results being presented as in Figure 2C. (**A**) Burst size increased with MP depolarization (**B**), Intra-burst frequency increased with MP depolarization (**C**), Relative AP threshold became more negative with MP depolarization (**D**) AHP amplitude increased with MP depolarization (**E**) AHP repolarization speed increased with MP depolarization. Data are averaged per burst and shown for each cell. The first AP in the burst was not taken into account. **F:** Cells with the more depolarized Vrest showed a greater effect of AP-prepotential (= ‘AHP amplitude’ for within burst APs) on AP threshold. Note that slopes of the correlation between relative AP threshold and AP-prepotential were bigger at more depolarized Vrest. **G:** Model quantification of the effect of membrane hyperpolarization and Nav channels availability on AP threshold, showing an amplification of the AP threshold shift produced by membrane hyperpolarization when starting Vrest is more depolarized (note the change of the slope of the correlation between AP threshold and pre-spike hyperpolarization). **H:** Curve depicting the modification of AP threshold model as a function of the quantity of Nav conductance available before the spike. Note that the same 10 mV membrane hyperpolarization produced a stronger modification of AP threshold when Vrest is more depolarized

### The evolution of a late slow component of the AHP contributed to burst termination

After taking an interest in the burst initiation mechanisms, let us now focus on mechanisms leading to the burst stop. A plausible mechanism that could account for burst termination would be an evolution along the burst that would return the AP threshold to values above the Vrest. To evaluate for this possibility, we plotted the AP threshold evolution and compared the AP threshold modification, relative to Vrest, between the second and the last AP of the burst (same method as in figure 8). We found only a negligible depolarization of AP threshold (Fig 11A, increase of normalized threshold: mean 0.053 ± 0.027 mV, paired t-test: t = 3.83, p < 0.001, N = 1276, ES = 0.11) that however remained largely below or within the range of subthreshold oscillation of the MP. Such a change probably reflected the small decrease of absolute AHP amplitude and small slowing-down of AHP repolarization rate along the bursts: see Fig 11B for amplitude (from the 2^nd^ to last burst AP: mean: 0.46 ± 0.04 mV, paired t-test: t = 23.5, p < 0.001, N = 1276, ES = 0.65, change occurring only between 2^nd^ and 3^rd^ Aps) and Fig 11C for repolarization rate (from 2^nd^ to last AP: −132 ± 9 mV/ms, paired t-test: t=−27.8, p < 0.001, N = 1276, ES = −0.77, change occurring mainly during the first 10 APs, the average slope being then constant).

**Fig 11:**
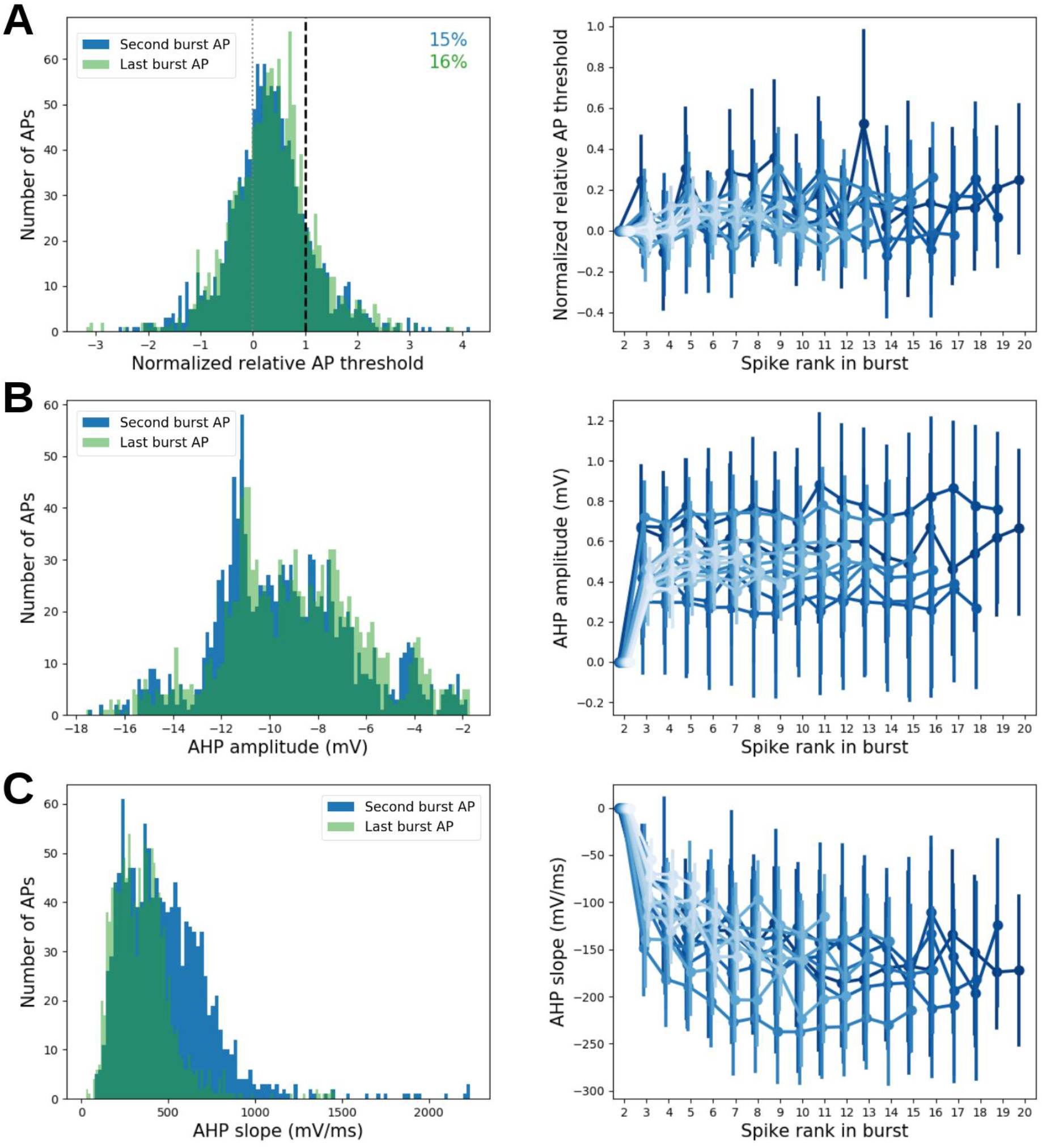
Evolution of MP dynamics during bursts. **A:** histograms of normalized AP threshold (linear interpolation between 0 and 1 which were respectively the resting potential and the maximum amplitude of subthreshold fluctuations) for second AP (blue) and last AP (green) in bursts (left panel). Right panel shows the evolution as a function of AP rank in burst of the normalized AP threshold. To compare across bursts, data wer shifted and aligned at 0 for the second AP threshold. APs were grouped per burst size (gradient of blue colours). Error bars represent 5-95% confidence interval of the mean. We observed a clear but small increase of normalized AP threshold during burst. **B:** Same graphs as A but for AHP amplitude (preceding the AP). Right panel clearly shows that there was a small decrease of absolute AHP amplitude from 2^nd^ to 3^rd^ AP in burst, but after 3^rd^ AP in burst the average AHP amplitude is constant. **C:** Same graphs as A but for AHP slope (preceding the AP). There was a clear decrease of AHP slope during the first 10 APs in burst.

To clarify the burst termination mechanisms we used of a linear model (using Vrest, AHP amplitude, AHP slope and AHP duration as parameters, see Methods for details) which predicts the dynamics of putative AP threshold after the last AP of the burst (see examples in Fig 12A and 12B, dashed lines). This analysis showed that for 89% of the bursts that the MP that followed the last AP did not overcome the putative thresholds (Fig 12D). Such an effect cannot be explained by the threshold value variation which remained, at a time interval equal to the last inter-spike interval, similar to that of the last AP of the burst (see Fig 12C, difference of potential between last AP threshold and predicted threshold: 0.09mV, SD: 0.68mV, N = 1532 bursts). The failure of MP to overcome the AP threshold appeared to be due to the appearance of a late and slow component in the AHP, after the early fast component characterising the intraburst AHPs repolarizing phase. This new AHP component could be fitted with a slow exponential (with a time constant estimated for all cells (mean: 171.66ms, SD: 88.76ms, see Fig 12E and F, see also figure 2C in Balu and Strowbridge, 2004). This component kept the MP more hyperpolarized than during the intraburst AHPs, preventing the MP to reach the AP threshold. The results suggested a scenario according to which the AP threshold could be reached only during the early fast component of the repolarization phase of the AHP, the appearance of the late slow repolarization precluding the MP to reach a threshold that would have thereafter rapidly moved to pre-burst values.

**Fig 12:**
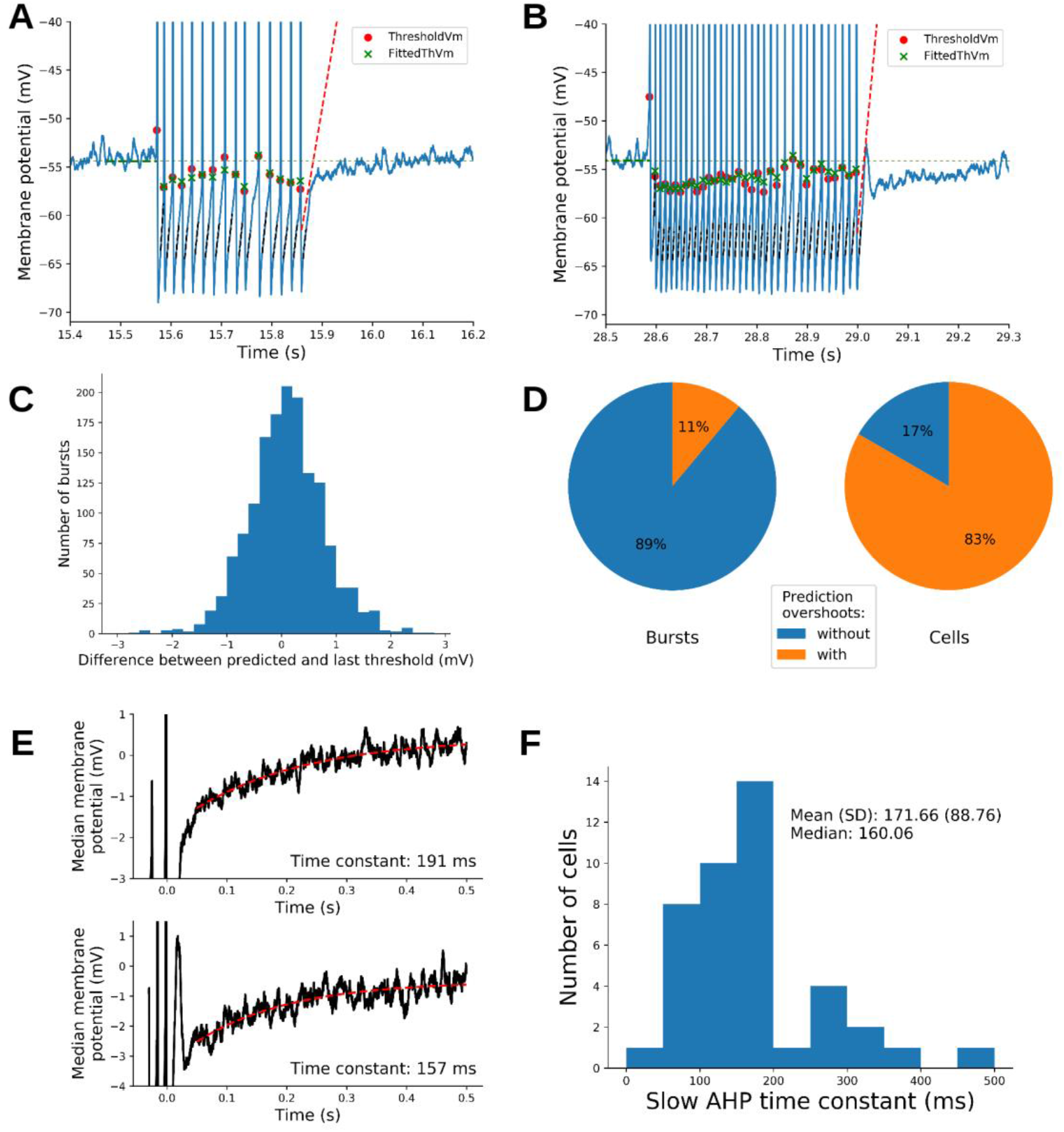
Prediction of post-burst AP threshold and slow AHP component. **A-B:** Examples of bursts with detected AP threshold (red dots) and their model fits (green cross). Dashed red lines show the predicted threshold as function of time elapsed after the beginning of the last AHP. We noted that bursts could be followed by post-burst rebound (**B**) or not (**A**). In both cases the MP stayed below the predicted AP threshold accounting for the end of the burst, and a sudden decay of MP repolarization speed was observed. See text and material and methods for model details. **C:** Distribution of the differences between predicted AP threshold (at the same ISI as the last burst ISI) and the threshold of the last AP in the burst. **D:** As shown in A, we could detect when signals at burst end was going above the predicted threshold (overshoot) without AP firing. Left pie chart shows the proportion of burst with an overshoot (pooled across cells). Right pie chart shows the proportion of cells with at least one overshoot among their bursts. **E:** Median traces of the 500 ms following the last AHP peak with exponential fits of the slow AHP component (calculated from 50ms to 500ms following AHP peak). Upper and lower panels correspond respectively to cells shown in A and B. **F:** Distribution of exponential time constants from fits as shown in **E** for all cells computed here for 42 cells (see Methods for details)

Thus, the arrest of burst discharge seems to be attributable to the apparition of a slow component during the AHP repolarizing phase In order to investigate whether this slow component developed along the burst, we performed the experiment depicted in figure 13. Here MCs were slightly hyperpolarized with a steady current injection preventing spontaneous firing and, APs were evoked by short (3 ms) depolarizing current steps. Five bursts of 1, 2, 4, 8 and 16 AP respectively were generated and, the last AHP parameters were compared (Fig 13A). As shown in figure 13B-C of AHP amplitude and area increased with the increase of the number of APs (repeated measures ANOVA p<0.001, N=11). Interestingly, the slow AHP component appeared at more and more hyperpolarized MP as the number of APs increased(repeated measures ANOVA p<0.001, N=11. Fig 13E). This result suggests that, along the burst, the slow AHP component would appear at more and more hyperpolarized potentials, thus decreasing the probability that the fast AHP repolarization could overcome the AP threshold The slow late AHP component is reminiscent of the slow inactivating K^+^ current that was previously observed in MC following membrane hyperpolarization and, that was suggested to be produced by the recovery from inactivation of I_A_ current (See figure 5 of Balu end Strowbridge 2007). The hypothesis that the slow AHP component was due to I_A_ was further supported by a computational model of MC showing the increase of this current during the AP burst (Rubin & Cleland, 2006). Indeed, application of 4AP (3 mM) prevented the evolution of the AHP during the evoked burst (AHP area: 4AP effect p=0.005; interaction p<0.001; AHP amplitude: 4AP effect p=0.002; interaction p<0.001; Fig 13D) as well that of Vm toward hyperpolarized values, at which the slow AHP component appeared (4AP effect p=0.004; interaction p<0.001; Fig 13E). Note that 4AP reduced both the early-fast and the late slow AHP. The hypothesis that the slow component would involve the activation of I_A_ current is further supported by our toy model (Fig 13 F). In fact, in the simple model involving only Nav and Kdr channels, there was no development of the slow AHP component as the number of APs increased (Fig 13 F, no I_A_). However, the simple implementation of I_A_ current to this model reproduced the development of the slow AHP with the number of APs found in the experiments (Fig 13 F, I_A_ and I_A_ modified). The conductance of I_A_ was either directly taken from (Rubin & Cleland, 2006) or modified to get biophysics closer to previously published I_A_ biophysics (Amendola *et al.,* 2012) (see methods). Noteworthy is that the I_A_ current dynamic has a maximal inactivation at the AHP peak and then a reactivation during the repolarization phase of the fast AHP. Altogether these data suggested that the burst is stopped by the build-up of 4-AP-dependent IA-like current that, by slowing down the AHP, brings the AP threshold to values that cannot be reached during AHP repolarization.

**Fig 13:**
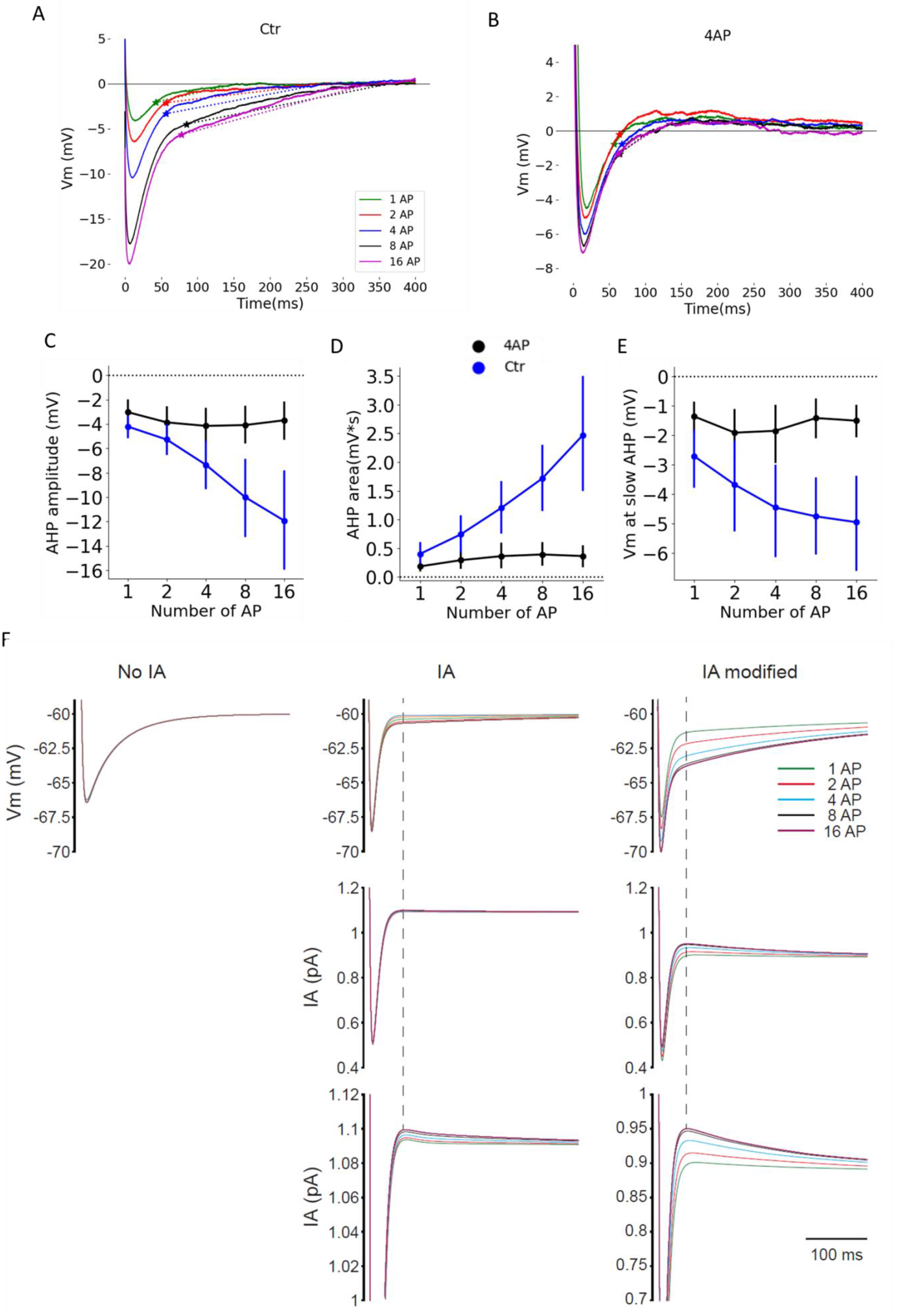
The AHP evolution with the number of APs was mainly due to 4AP sensitive component. **A)** Superposition of the last AHP of evoked bursts constituted of different number of APs. Asterisks indicate the onset of the AHP slow component i.e. at which the “Vm at slow AHP” was calculated. **B)** Same neuron as A but in presence of 4AP (3 mM). **C)** Modification of AHP, as a function of the number of APs in the bursts, in control condition and in presence of 4AP. **D)** Modification of AHP area, as a function of the number of APs in the bursts, in control condition and in presence of 4AP. **E)** Modification of the MP at which the slow component appeared, as a function of the number of APs in the bursts, in control condition and in presence of 4AP. **F)** Reduced toy model showing the superposition of the AHPs at the end of bursts constituted of different number of APs. Top, with only Nav and Kdr channels, there was no evolution of the AHP by contrast, implementation of I_A_ channels reproduced the evolution of the slow AHP observed in experimental data. Middle and bottom, evolution of I_A_ current during the AHP at different magnifications. IA, channels biophysics from (Rubin and Cleland, 2006); I_A_ modified, channels biophysics closer to Amendola and colleagues(2012). Error bars represent 95% Confidence Interval. N = 11 MCs.

## Discussion

Our study provides new insights into the understanding of the intrinsic cellular mechanisms responsible for the genesis of firing activity in MCs. More precisely, we have shown that AHP plays a key role in this genesis since changes in its characteristics (duration, amplitude, kinetics) can both trigger and stop the burst generation while also determine the bursting properties. The experimental results presented in this report have been synthetized to build the model depicted in figure 14. According to the model, the firing of MC is triggered by a modification of the AP threshold that dynamically changes as a function of the MP trajectory. Due to the relatively depolarized Vrest of MCs, a part of the voltage dependent Na^+^ channels are inactivated. The availability of these channels may increase under the influence of the intrinsic oscillatory activity and synaptic inputs received by MCs; both of which can shift MP towards more hyperpolarized value, bringing the AP threshold within MP subthreshold fluctuations, thus, facilitating the firing. Once the first AP is generated, the following fast AHP brings the threshold below the Vrest, or within the Vrest noise, acting in this way as a regenerative mechanism that will produce the burst.

**Fig 14:**
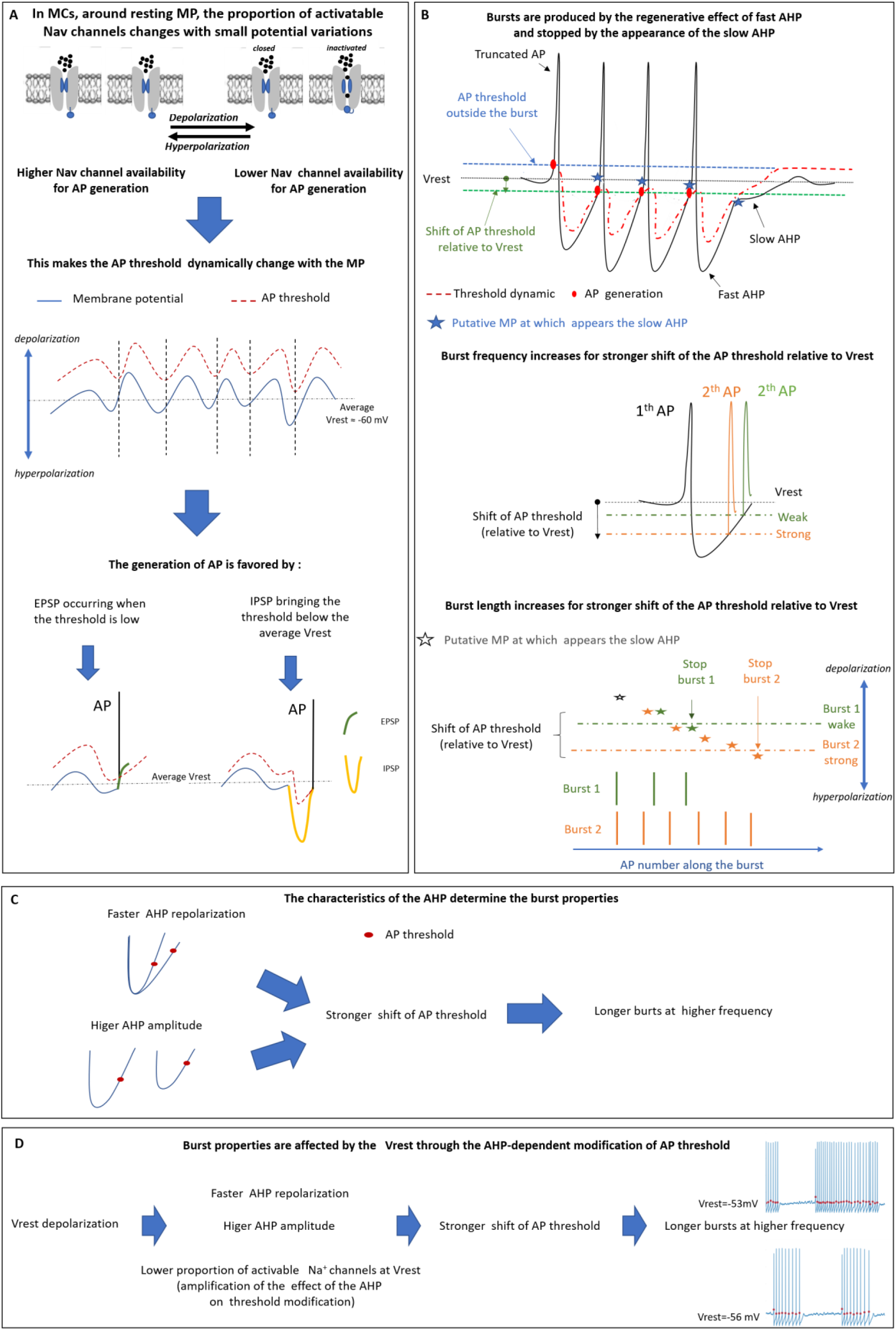
Model of the intrinsic mechanisms accounting for MC firing. **A)** Change of deinactivated/inactivated state of Nav channels make the AP threshold dependent on MP fluctuations. Due to the deinactivation kinetics threshold fluctuation (red dashed line) are shifted compared to MP fluctuations (blue line). The generation of the AP is favoured by excitatory inputs occurring just after the negative phase of MP fluctuations or when the repolarization phase of inhibitory inputs is rapid enough to overcome the shifted threshold before the latter goes back to the pre-ipsp values. **B)** The fast AHP brings the AP threshold below the Vrest and its repolarization rate is rapid enough to overcome the modified threshold (shift of AP threshold), acting in this way as a regenerative mechanism that sustains the burst. The late and slow components of the AHP develop gradually along successive APs. When the MP, at which the slow component should appear, is above the shift of AP threshold, the manifestation of the slow component is bypassed by the AP generation Indeed, the slow AHP can manifest itself only at MP more negative than the AP threshold shift. In such condition, the MP repolarization during the slow component is not fast enough to overcome the AP threshold, which then reverts to the pre-burst value, and the burst stops. For a stronger shift, the AP threshold is reached earlier, during the repolarization phase of the fast AHP, and the intra-burst frequency is higher. For stronger shift of AP threshold, a higher number of APs is necessary so that the slow AHP could manifest itself and thus, the burst duration increases. **C)** The threshold shift is higher when the fast AHP is larger and faster. In such conditions, the bursts are longer and have higher intra-burst frequency. **D)** The AP threshold shift is higher when the Vrest of the MC is more depolarized, because of modifications of AHP properties and partial inactivation of Nav channels. As a consequence, MP depolarization makes bursts longer and with higher intra-burst frequency.

The burst termination is ensured by a slow, 4AP dependent, AHP component that progressively develops along the consecutive APs, and is hypothesized to involve I_A_ current. This component slows down the AHP repolarization phase, thus increases the inactivation rate of Na^+^ channels and moves back the AP threshold to values that cannot be overcome by MP repolarization.

The intraburst properties (frequency, length) are determined by the magnitude of the modification of AP threshold relative to the Vrest. The larger the modifications of AP threshold, the longer are the bursts (in term of number of APs) and the higher are intraburst firing frequencies. The firing frequency increases because the AP threshold is reached faster during the AHP repolarization phase, especially when AP threshold shift towards hyperpolarizing values is associated with faster AHP repolarization. Burst length increases because the slow component needs more APs in order to manifest itself at more negative MPs than the AP threshold. Indeed, because the AP-threshold shift during the burst is driven by the AHP, the burst properties are determined by the AHP features. In particular, the number of APs and firing frequency increase, when AHP amplitude and repolarization rate increase. The model predicts the burst length and intra-burst frequency increases that are observed upon membrane depolarization. In this condition the AHP amplitude increases, possibly due to an increase of K^+^ driving force and, the AHP repolarization becomes faster, by a yet unknown mechanism. Moreover, upon membrane depolarization the number of inactivated Nav channels augments. As a consequence, the recovery from inactivation of Nav channels produced by the AHP entails a greater shift of AP threshold relative to Vrest. In fact, our toy model showed that a decrease of the number of Nav channels available entails an increase of the hyperpolarization-induced AP-threshold shift (Figure 10 G-H).

Altogether, the heterogeneity of the firing properties observed among the different MCs would be therefore mainly due to differences in shape of AHPs, these differences being based on the relative importance of its two components (amplitude and slope), themselves directly depending on MP value history.

However, our model does not predict the minority of intraburst APs for which the threshold was clearly above the Vrest (Fig 8C-D). A mechanism that could account for these events is a rebound depolarization that was frequently observed at the end of the AHP (see Figures 8B and 12B for some examples). Obviously when it could be detected, the rebound depolarization did not overcome the AP threshold, but a great variability in the rebound amplitudes were observed, supporting the above hypothesis. With regard to the post AHP rebound depolarization, it could be due to the activation of persistent Na^+^ current (Balu *et al.,* 2007).

Not only depicting our own results, our model fits with others reported in the literature. As an example, tufted cells of the OB, displaying higher amplitude and faster repolarization AHP than MCs show, as our model would predict, a more sustained bursting activity; namely longer and higher frequency bursts (Burton & Urban, 2014). Similar covariation between AHP and burst properties have been also reported in MCs during postnatal development (Yu *et al.,* 2015). Our model predicts that these covariations are the consequence of AHP dependent modifications of the AP threshold shifts As for burst termination, in agreement with our hypothesis on the involvement of I_A_ current, the application of 4AP, has been reported to transform MC bursting activity into a continuous firing one (Balu *et al.*, 2004). MCs present a dendritic recurrent synaptic transmission that is characterized by a glutamatergic auto excitation (Aroniadou-Anderjaska *et al.,* 1999; Friedman & Strowbridge, 2000; Salin *et al.,* 2001) and a feed-back inhibition that follows the activation of granular cells (Isaacson & Strowbridge, 1998; Schoppa *et al.,* 1998). We have recently shown that the main effect of recurrent synaptic transmission is to shape the AHP of MC (Duménieu *et al.,* 2015). In particular, the recurrent inhibition increases the amplitude of the AHP without affecting the medium or the late AHP, while the recurrent excitation reduces both the amplitude and the medium component of the AHP (see Duménieu et al., 2015 figure 5D). Based on the present results we can therefore speculate that recurrent synaptic inhibition would favour long bursts at higher firing frequency; by contrast, the functional role of recurrent excitation is less predictable. Indeed, while the reduction of the AHP amplitude would favour short bursts at low frequency, we do not know whether and, to what extent, the reduction of medium component would affect the velocity of the repolarization phase of the AHP and, further analysis on this aspect is needed to elucidate the role of recurrent excitation on MC firing properties.

Let us focus on the interpretation of some of our experimental results which are less immediate and that requires further discussion: in particular, the contribution of I_A_ in the AHP course (Fig 13) and the role of the recovery of sodium channel in the modification of AP threshold (Fig 5). As evidenced from figure 12, an increase in AHP amplitude was observed with an increase of the number of evoked consecutive APs. This appeared as mainly based on the increase of 4AP-dependent AHP-component. Noteworthy was that such AHP amplitude increase was not observed during spontaneous burst (Fig 11B). One possible explanation is that I_A_ would develop earlier when the APs are evoked by experimental depolarizing steps having a long duration (3 ms), relatively to the AP half width (~ 0.7 ms). Moreover, the depolarizing step could mask the veritable starting point of the AHP, giving the impression that I_A_ component was already present since its beginning. It is therefore plausible, that during spontaneous firing activity I_A_ appears early enough to affect the medium AHP, and therefore slowing down the AHP repolarization during the burst (Fig 11C), but, too late to affect the AHP amplitude. The hypothesis of the involvement of the recovery from inactivation of Nav, in threshold modification, would predict an increase of the effect of hyperpolarization when the global availability of Nav channels was reduced (fig 5E). This prediction was confirmed by the application of TTX at 20 nM but not when TTX at 10 nM was used. However, 10 nM already reduced the AP threshold produced by the depolarization steps, suggesting that it reduced the availability of Nav (Fig 4B). This absence of TTX effect at 10 nM is unlikely a consequence of sampling variability (i.e. a false negative result) since Bayesian analysis supports an actual absence of the effect (BF10<1/3; Keysers et al 2020). Moreover, when selecting only neurons for which both 10 and 20 nM TTX were applied, 20 nM reliably increases the effect of the MP hyperpolarization on AP threshold while 10 nM did not (data not shown). A plausible interpretation of this result is that the recovery from inactivation - produced by membrane hyperpolarization- may involve only one/some of the different sub-types of Nav channels which would be low sensitive TTX ones and would be not yet blocked at 10 nM. Further investigations are needed to confirm such hypothesis, for example, by using selective antagonists of the different Nav subtypes.

### Conclusions

Our results show that in MCs, the AHP characteristics play a pivotal role in determining the properties of MCs’ firing activity. AHP characteristics have been shown to be modified by the membrane potential (present work), neuromodulation (Wu *et al.,* 2002; Brosh *et al.,* 2006), postnatal development (Duménieu *et al.,* 2015; Yu *et al.,* 2015), learning (Duménieu *et al.,* 2015; Reuveni & Barkai, 2018) and recurrent synaptic transmission (Duménieu *et al.,* 2015). Altogether, AHP appears to be a key target which, thank to highly scalable characteristics, could modulate OB processing according to many parameters such as physiological state (reproduction period, food need, etc.), memory and experience.

## Acknowledgment

This work was supported by the CNRS, Inserm, and Lyon 1 University.

We are grateful to Marco Canepari for reading the manuscript and suggestions.

